# Unraveling Neural Complexity: Exploring Brain Entropy to Yield Mechanistic Insight in Neuromodulation Therapies for Tobacco Use Disorder

**DOI:** 10.1101/2023.09.12.557465

**Authors:** Timothy Jordan, Michael R. Apostol, Jason Nomi, Nicole Petersen

## Abstract

Neuromodulation therapies, such as repetitive transcranial magnetic stimulation (rTMS), have shown promise as treatments for tobacco use disorder (TUD). However, the underlying mechanisms of these therapies remain unclear, which may hamper optimization and personalization efforts. In this study, we investigated alteration of brain entropy as a potential mechanism underlying the neural effects of noninvasive brain stimulation by rTMS in people with TUD. We employed sample entropy (SampEn) to quantify the complexity and predictability of brain activity measured using resting-state fMRI data. Our study design included a randomized single-blind study with 42 participants who underwent 2 data collection sessions. During each session, participants received high-frequency (10Hz) stimulation to the dorsolateral prefrontal cortex (dlPFC) or a control region (visual cortex), and resting-state fMRI scans were acquired before and after rTMS. Our findings revealed that individuals who smoke exhibited higher baseline SampEn throughout the brain as compared to previously-published SampEn measurements in control participants. Furthermore, high-frequency rTMS to the dlPFC but not the control region reduced SampEn in the insula and dlPFC, regions implicated in TUD, and also reduced self-reported cigarette craving. These results suggest that brain entropy may serve as a potential biomarker for effects of rTMS, and provide insight into the neural mechanisms underlying rTMS effects on smoking cessation. Our study contributes to the growing understanding of brain-based interventions for TUD by highlighting the relevance of brain entropy in characterizing neural activity patterns associated with smoking. The observed reductions in entropy following dlPFC-targeted rTMS suggest a potential mechanism for the therapeutic effects of this intervention. These findings support the use of neuroimaging techniques to investigate the use of neuromodulation therapies for TUD.

## 1. Introduction

Brain-based neuromodulation therapies, such as repetitive transcranial magnetic stimulation (rTMS), are emerging as a new class of treatments for substance use disorders. A notable milestone has been regulatory (FDA) approval of rTMS to treat tobacco use disorder (TUD). Approval was granted on the basis of a multicenter, double-blind, randomized controlled trial finding higher smoking cessation rates in individuals who received active vs. sham rTMS (Zangen et al., 2021). This important finding capitalizes on many previous studies demonstrating that excitatory rTMS to the left dorsolateral prefrontal cortex (dlPFC) increases smoking abstinence rates relative to sham (Dinur-Klein et al., 2014; X. Li et al., 2020), lowers the rates of relapse to smoking (Sheffer et al., 2018), reduces cigarette craving (X. Li et al., 2013; Pripfl et al., 2014), and reduces the number of cigarettes smoked (Abdelrahman et al., 2021; Amiaz et al., 2009; Huang et al., 2016; X. Li et al., 2020; Prikryl et al., 2014). This body of work has led to clinical recommendations advocating for the use of rTMS as a smoking cessation treatment (Young et al., 2021), and suggests that this promising new brain-based therapeutic can be informed by advances in neuroimaging that shed light on the neural circuitry alterations associated with TUD.

Noninvasive neuromodulation treatments have been designed to capitalize on findings linking the insula to smoking cessation by attempting to stimulate the insula. The neural basis of TUD has been strongly linked to the insula via lesion studies, finding that lesions to the insula per se (Naqvi et al., 2007) and a broader network involving the insula (Joutsa et al., 2022) are associated with higher rates of smoking cessation compared to lesions involving other brain regions. Structural MRI studies also link the insula and other associated brain regions to TUD. People who smoke have smaller gray matter volumes in areas of the prefrontal cortex and anterior cingulate (Brody et al., 2004). More cigarette exposure is associated with thinner insular cortex (Morales et al., 2014), and in women, thinner insular cortex is associated with more cigarette craving (Perez Diaz et al., 2021). Both meta-analysis (Hill-Bowen et al., 2022) and mega-analysis (Mackey et al., 2019) have reported smaller amounts of gray matter in both the medial prefrontal cortex and insula in individuals with varying substance use disorders, with a smoking-specific effect (i.e., not found in individuals with other kinds of substance use disorders) of smaller volumes in the posterior cingulate cortex; Hill-Bowen et al., 2022).

Resting-state functional connectivity studies have found that both the insula specifically, and also large-scale network dynamics involving the insula, are implicated in both acute and chronic nicotine use. People who smoke have lower overall functional connectivity in the brain (Cheng et al., 2019), which has also been shown specifically within the executive control and default mode networks (Weiland et al., 2015). Functional connectivity features can be used in machine learning to distinguish between people who do and do not smoke (Wetherill et al., 2019). A triple-network model describing the relationship between the salience, default mode, and executive control networks (Fedota & Stein, 2015) suggests that the relationship between these three networks responds dynamically to nicotine use: smoking increased coupling between the left executive control network and salience network, and decreased the anticorrelation between the default mode network and salience network (Lerman et al., 2014).

Resting-state functional connectivity analyses have also specifically linked the insula to cigarette craving, withdrawal, and relapse. The magnitude of withdrawal correlates positively with the strength of connectivity between the right ventral anterior insula and dorsal anterior cingulate cortex (Ghahremani et al., 2021), both hubs of the midcingulo-insular network (also referred to as the salience, cingulo-opercular, or ventral attention network; Uddin et al., 2019). Stronger connectivity between the insula and cortex surrounding the central gyrus is associated with better cessation outcomes (less relapse) (Addicott et al., 2015); similarly, stronger connectivity between the ventral striatum and a network including the insula is associated with better cessation outcomes(Sweitzer et al., 2016).

The insula is anatomically located underneath the cortical surface, rendering it inaccessible to conventional rTMS devices, but a small proof-of-concept study(Moeller et al., 2022) and electric field modeling work have suggested that deep rTMS machines such as the BrainsWay® H4 coil can penetrate deeply enough to reach the insula (Fiocchi et al., 2018). However, network connectivity may provide an alternate route to stimulate the insula and other brain regions involved in TUD(X. Li et al., 2017) using conventional rTMS devices.

Though stimulating left dlPFC shows promise for smoking cessation treatments, the exact mechanism that it works through remains unclear. Although rTMS undoubtedly produces salutary behavioral effects, and in some cases produces widespread changes in functional connectivity that can extend outside the stimulated network(Beynel et al., 2020), rTMS to most brain regions does not appear to change BOLD signaling at the stimulation site(Rafiei & Rahnev, 2022). Therefore, the mechanism by which rTMS causes changes in the stimulated region and other associated regions to yield changes in behavior remains ambiguous. Developing, optimizing, and personalizing these techniques may be improved by a more comprehensive understanding of normal brain function, the brain dysfunction associated with substance use, and the brain function that underlies responses to neuromodulation. Measuring brain entropy is an emerging approach that offers the potential to extend existing knowledge of brain features associated with substance use disorders.

Brain entropy quantifies the complexity and unpredictability of brain activity – as opposed to measures such as Pearson correlations, which measure the association between two brain regions, or standard deviation, which assesses variability. Sample entropy (SampEn) is an approach developed in the context of information theory that has recently been applied to understand the structure of neural time series data. By measuring the similarity between two components (subsequences) of a time series, SampEn quantifies regularities and irregularities, and thereby provides information about the complexity and predictability of the time series signal. A validation study has shown that SampEn can be accurately determined for fMRI data on both simulated and actual datasets(Z. Wang et al., 2014). Higher SampEn values reflect time series that are more complex and therefore less predictable, and conversely, lower SampEn values reflect time series that are less complex and therefore more predictable. SampEn is less sensitive to noise and abbreviated data sets than other forms of entropy (e.g., Shannon entropy and Approximate Entropy), rendering it a good candidate for analysis of fMRI time series data(Richman & Moorman, 2000; Z. Wang et al., 2014; Yentes et al., 2013).

Previous work has suggested that SampEn is both altered by rTMS(Song, Chang, Zhang, Peng, et al., 2019) and is also different in neuropsychiatric populations compared with healthy individuals, although the direction of the effect depends on the population studied. Individuals with Attention Deficit Hyperactivity Disorder (ADHD) have lower frontal and occipital entropy compared with controls, and symptom severity correlates negatively with entropy levels(Sokunbi et al., 2013). Similarly, lower SampEn measurements have been observed in patients with Alzheimer’s disease(B. Wang et al., 2017). Notably, machine-learning classifiers that use SampEn to distinguish patients from controls outperform those relying on standard correlation-based measurements(Wu et al., 2021). Likewise SampEn has been shown to correlate with fractional amplitude of low-frequency fluctuation (fALFF) measurements (Song, Chang, Zhang, Ge, et al., 2019; Zhang et al., 2021), network coherence frequency ranges (D. J. J. Wang et al., 2018) and power spectrum measures (Bruce et al., 2009) showing that changes in SampEn captures a diverse set of neural mechanisms, which is a desired feature in a biomarker.

In contrast to the relatively lower SampEn observed in individuals with ADHD and Alzheimer’s Disease, higher SampEn has been observed in people who smoke(Z. Li et al., 2016), and a small pilot study suggested that noninvasive neuromodulation using repetitive transcranial magnetic stimulation (rTMS) can reduce both entropy and cigarette craving in these individuals(Song, Chang, Zhang, Peng, et al., 2019). Demonstrating that rTMS can influence entropy in people who smoke could fill this gap in knowledge about rTMS mechanisms, so we sought to test the hypotheses that (1) high-frequency rTMS to the dlPFC would reduce SampEn in the dlPFC and insular cortex, and (2) greater reductions in SampEn in these regions would correspond with greater reductions in craving.

## 2. Methods & Materials

### 2.1 Participants

To test the above hypotheses, data were collected from 42 participants. All participants were recruited from the greater Los Angeles community. Study procedures were approved by the UCLA IRB and written, informed consent was obtained from all participants for being included in the study. Initial eligibility assessments were made by telephone, and participants who met criteria according to their self-report were scheduled for further in-person eligibility screening, which included baseline neuroimaging measurements (see below).

To be included, participants were required to be right-handed, between the ages of 18 to 45, smoking on average 5 or more cigarettes per day, and not seeking or receiving treatment for smoking cessation. Participants were excluded from participation if they were left-handed, met criteria for any other substance use disorder, met criteria for other psychiatric conditions as assessed by the Mini International Neuropsychiatric Interview version 7.0.2 (Sheehan et al., 1998); tested positive for other substances of abuse by urinalysis or breathalyzer; if they reported or tested positive for pregnancy; or if they were determined to have safety contraindications for rTMS or MRI, including non-removable metal implants or any factor that could lower the seizure threshold. **Table 1** shows the demographic characteristics of the participants included in this study.

**Table 1:** Demographic Table. Study project demographics with number of participants included, N, the mean age of the participants in years, the number of males and females, and the average duration of abstinence prior to their test session for each stimulation site.

| Demographic Table |  |  |
| --- | --- | --- |
| N |  | 45 |
| Years of age (mean ± SEM) | | 33 ± 7 |
| Years of Smoking (mean ± SD) | | 15.2 ± 1.24 |
| Sex | M | 33 (73.3%) |
|  | F | 12 (26.7%) |
| Hours of abstinence | dIPFC | 16±1 |
| | v5 | 16±0.95 |

### 2.2 Study Design

Participants who remained eligible after in-person assessments were scheduled for data collection sessions. Each session was identical except for the region stimulated (left dlPFC or visual cortex [v5]). The order of each session type was randomized and counterbalanced, and participants were instructed to remain abstinent from smoking for >12 hours before their data collection sessions.

Upon arriving in the laboratory for each data collection session, urine samples were collected to confirm abstinence from illicit substances, a breathalyzer was administered to confirm abstinence from alcohol, and vital signs were obtained. Expired carbon monoxide was measured to confirm >12 h abstinence from cigarette smoking.

To assess baseline withdrawal and craving, participants completed the Shiffman-Jarvik Withdrawal Questionnaire (S. M. Shiffman & Jarvik, 1976) and Urge to Smoke scales (Jarvik et al., 2000) via self-report. Baseline resting-state functional images (see below for sequence details) were collected. On the first stimulation day only, the participant’s active motor threshold was measured and recorded. On each stimulation day, neuronavigation was used to position the rTMS coil. rTMS was delivered (see details below), followed immediately by a post-rTMS resting-state neuroimaging scan, then post-rTMS self-report of craving and withdrawal measurements. A CONSORT diagram of the study can be found in the supplementary materials. **Figure 1** shows a CONSORT diagram of the study.

**Figure 1.**
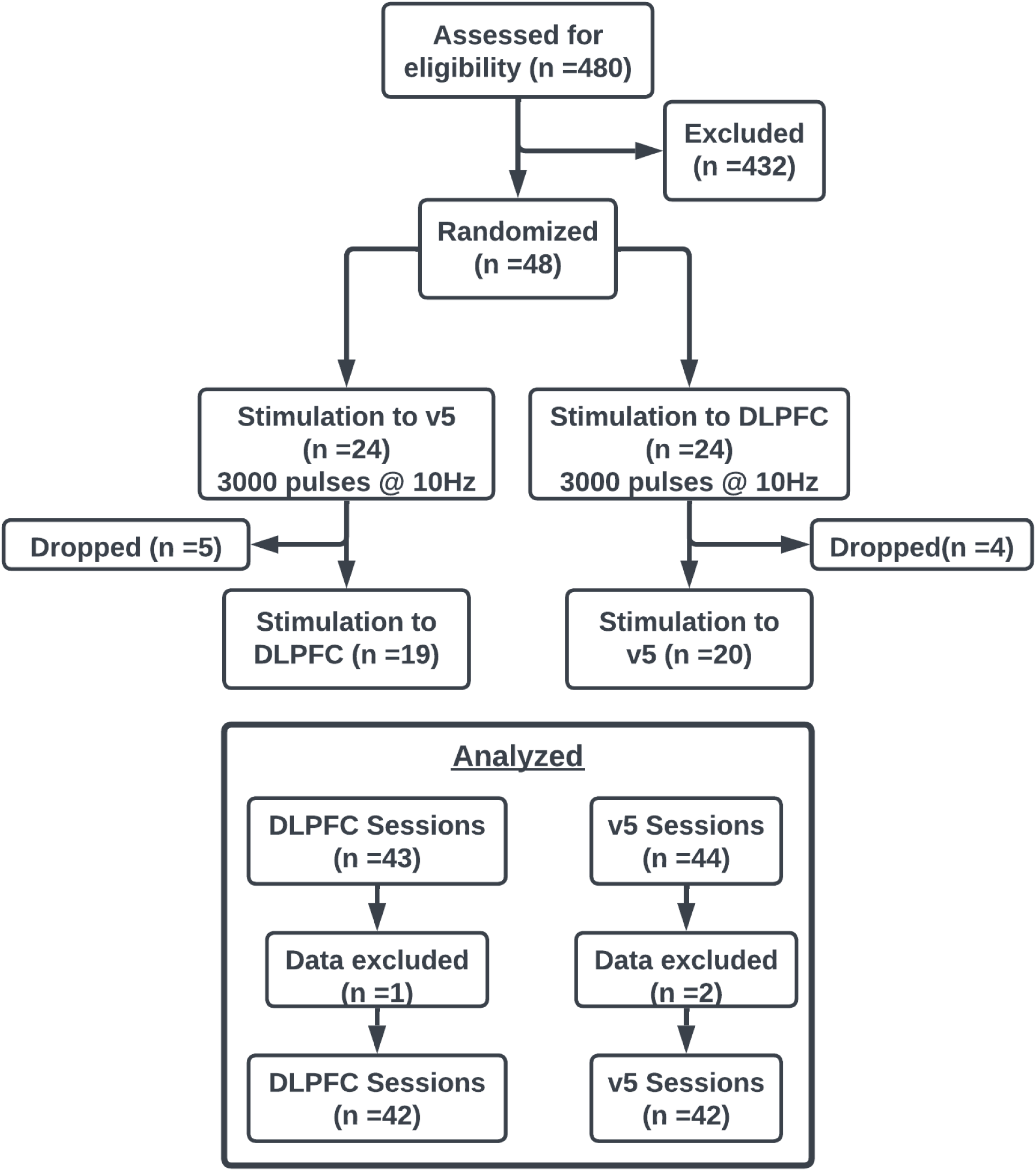
CONSORT Diagram. Our study design was a randomized one-way blind study. Eligible participants were randomized to either dlPFC first and v5 second, or vice versa. Participant data was not excluded if they only completed one session. Three sessions of data were excluded due to corrupted data or normalization errors.

### 2.3 Behavioral Data Collection

Participants were required to abstain from smoking for >12 hours before each testing day began to produce a state of acute withdrawal. Withdrawal is associated with a range of subjective experiences, including a heightened sense of craving as well as other somatic and affective symptoms. Although often linked together, withdrawal and craving are separate but related components of TUD (Baker et al., 2012; S. Shiffman et al., 2004). To access both variables, we used two questionnaires: Shiffman-Jarvik Withdrawal Questionnaire (SJWS,(S. M. Shiffman & Jarvik, 1976)) and Urge to Smoke scales (Jarvik et al., 2000). The SJWS assesses multiple domains of withdrawal, including a sub-scale specific to craving. For this study, we examined the SJWS overall score and the craving subscale scores in participants to capture both aspects in our participants. The craving sub-scale has two scores reported, the average and total scores, calculated from the questions specific to cigarette craving in the SJWS.

### 2.4 Brain Imaging Data Collection & Preprocessing

Whole-brain structural and functional MR imaging was conducted on a 3 Tesla Siemens Prisma Fit MRI scanner with a 32-channel head coil at the UCLA Staglin Center for Cognitive Neuroscience . A single T1-weighted structural scan (TE= 2.24ms ; TR= 2400ms; voxel resolution= 0.8 × 0.8 × 0.8 mm) was collected during the intake session as well as a 8 minute baseline T2*-weighted multi-band sequence resting state functional scan. Resting state functional scans (TE= 37ms; TR= 800ms; FoV = 208mm; Slice Thickness= 2mm; Number of Slices = 72, voxel resolution= 2 × 2 × 2 mm) were performed twice on test days (pre- and post-rTMS). Prior to all resting state functional scans, two spin echo fieldmaps were collected in opposite directions (AP and PA).

FMRI data processing was carried out using FEAT (FMRI Expert Analysis Tool) Version 6.00, part of FSL (FMRIB’s Software Library, www.fmrib.ox.ac.uk/fsl). The following pre-statistics processing was applied; motion correction using MCFLIRT(Jenkinson et al., 2002); B0 unwarping using boundary-based registration via FUGUE (Jenkinson, 2003, 2004); slice-timing correction using Fourier-space time-series phase-shifting; non-brain removal using BET (Smith, 2002); spatial smoothing using a Gaussian kernel of FWHM 4.0mm; grand-mean intensity normalization of the entire 4D dataset by a single multiplicative factor; highpass temporal filtering (Gaussian-weighted least-squares straight line fitting, with sigma=50.0s). ICA-based exploratory data analysis was carried out using MELODIC (Beckmann & Smith, 2004)[Beckmann 2004], in order to investigate the possible presence of unexpected artifacts or activation. ICA-FIX was trained on a set of 20 scans that were hand-classified into noise and non-noise components, with the scans randomly selected from 5 bins sorting scans by the amount of average motion present to have high and low motion data in the trained set. The component classification derived from the trained data was then used in ICA-FIX to classify noise and non-noise components from all subject data and non-aggressively remove the noise components. After denoising, ICA-FIX applied a high-pass filter to each subject’s data. Registration to high resolution structural and/or standard space images was carried out using FLIRT (Jenkinson et al., 2002; Jenkinson & Smith, 2001). Registration from high resolution structural to standard space was then further refined using FNIRT nonlinear registration (Andersson et al., 2007b, 2007a). Lastly, average time series were extracted from each subject’s data based on brain nodes specified by the atlas then detrended for cubic trends and finally z-score normalized.

### 2.5 Brain Atlas, dlPFC and Insula

For this study, we used the brain parcellation proposed by Van De Ville (Van De Ville et al., n.d.) to extract time series from all imaging data. Briefly, this parcellation includes a Schaefer 400 brain region cortical parcellation ((Schaefer et al., 2018), https://github.com/ThomasYeoLab/CBIG/tree/master/stable_projects/brain_parcellation/Schaefer2018_LocalGlobal) combined with 16 subcortical regions and 3 cerebellar regions from the HCP release for a total of 419 nodes. This study focused on changes in dlPFC and Insula; therefore, to determine which nodes correlated to those regions we used the Harvard-Oxford probability atlas and the dlPFC ROI mask obtained from Neurovault to determine which nodes were primarily in these regions. Insula was determined to overlay with nodes 35, 98 to 100, and 143 in the left hemisphere and nodes 234-236, 302-305 and 340 in the right hemisphere. Left dlPFC was determined to overlay with nodes 137 to 142.

### 2.6 Neuromodulation

#### 2.6.1 rTMS

TMS sessions were conducted using the Magstim Super Rapid2 Plus1 (MagStim, UK, https://www.magstim.com/row-en/) system equipped with a figure-8 coil. We stimulated two regions, dlPFC and v5, during separate sessions as shown in **Figure 2**. Stimulation to dlPFC was considered the active treatment region, while v5 served as a control region. Both stimulation sessions used the same stimulation sequence of 10 Hz stimulation for 60 trains, each train lasting 5 seconds and followed by 10 seconds of no stimulation, for a total of 3000 pulses over approximately 15 minutes.

**Figure 2.**
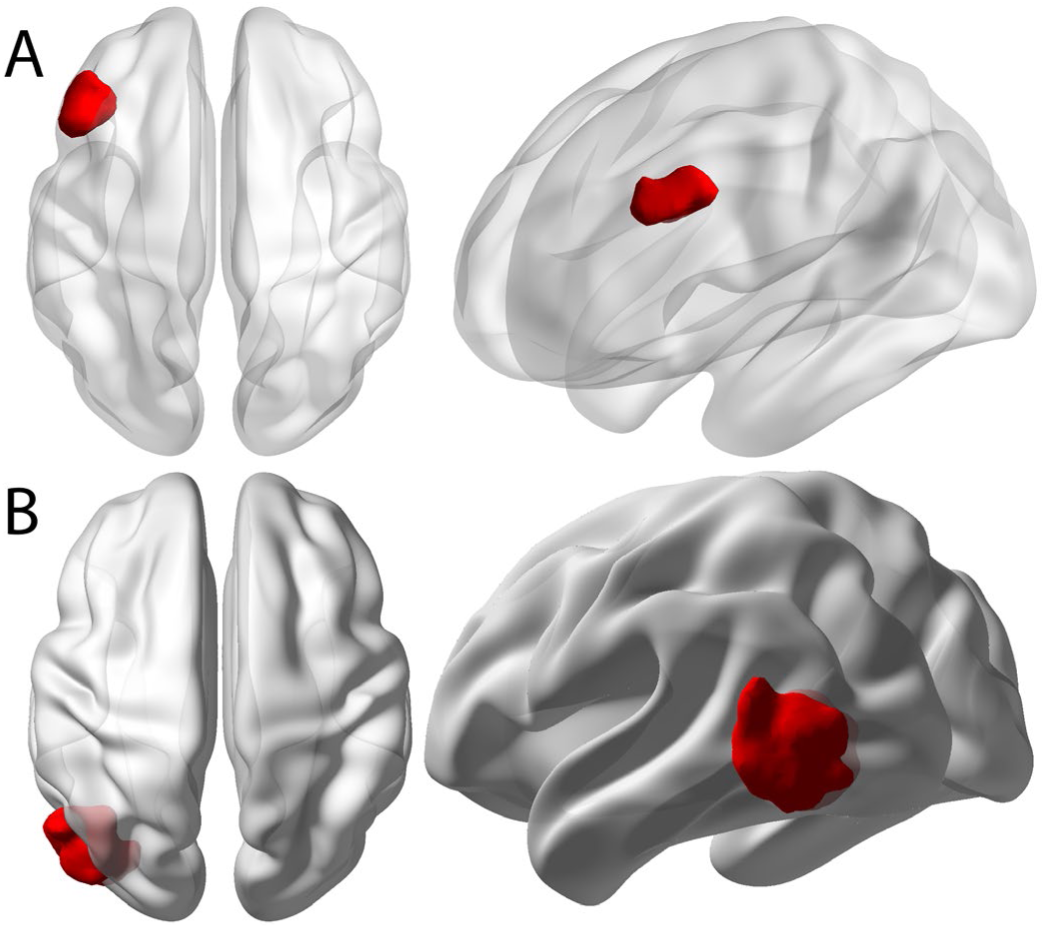
Stimulation Targets. A)The left dorsolateral prefrontal cortex (dlPFC) was used as the target region for this project. B) Counter to dlPFC, the left visual cortex (v5) was used as a control region

### 2.7 Analyses

#### 2.7.1 Brain Entropy Calculations & Analysis

Information entropy is the measure of randomness or uncertainty of a series of data without knowledge of the data series origin. One way to calculate this information entropy is called Sample Entropy. Given a data series, for example a time series extracted from a voxel or region of the brain, the process of Sample Entropy first divides the time series up into smaller vectors of length m. Next for each of the vectors it calculates the distance between the two vectors, with the requirement that they aren’t the same vector, i.e. i =/= j. If both values are less than the distance threshold (r), also can be called a noise filter as it determines the possibility of the pair, then the pairing is counted as a possible. The sum of all possible vector comparisons creates B(r), which is the probability that two sequences are similar for m points. This process is then repeated for vectors of m+1 size to determine the number of matches and sum those together and create A(r), which is the probability that two sequences are similar for m+1 points. Taking the ratio of the number of matches to the number of possibles, we find how much of the signal is uncertain/random. We then take the negative log of this ratio since information measurements are made on the logarithmic scale. The mathematical representation of this process is:

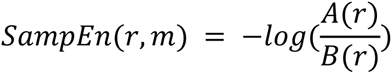

For this study, sample entropy of each node was calculated using the Brain Entropy Mapping Toolbox (BENtbx, (Z. Wang et al., 2014)). For the parameters, we set m = 3 and r = 0.3 based on a previous study examining the effects of these parameters on sample entropy (Yentes et al., 2013). Extracted mean node time series were organized into matrices with dimension timepoints by nodes, for the study this would generate a 588×419 matrix for 419 nodes each having 588 timepoints. The BENtbx would then take these matrices and calculate the entropy per node per participant.

## 3. Results

### 3.1 Brain Entropy Prior to rTMS

We examined the distribution of average SampEn across the brain at baseline before examining changes in SampEn due to brain stimulation. Using baseline (pre-stimulation) images, values for each node were collected from each participant and then averaged to determine the average resting SampEn for people who smoke and are in withdrawal. Likewise, all node values were averaged for each participant to determine their brain’s average SampEn. We then took the brain averages and compared them to each node’s values to determine if a region was significantly above or below the global average.

Considering both (1) the node average SampEn and (2) node average SampEn relative to brain average SampEn, we observed that gyral nodes had lower SampEn than the sulci nodes and the subcortical regions. ***Figure 3A*** shows the contrast between these areas of the brain. We also found that the majority of the outer cortical regions and the cerebellum were significantly below the global average (p_FDR_ < 0.05), and that the subcortical and orbital frontal regions were significantly above the cortex (p_FDR_ < 0.05). These observations and results complement and support previous findings by (Z. Wang et al., 2014). ***Figure 3B*** shows the regions found to be above (red) and below (blue) the average brain SampEn in people who smoke. Contrary to (Wang et al., 2014), who found that the range of SampEn values across the brain spanned from 0.44 to 0.608, in this study, we found that SampEn ranges across the brain spanning from 1 to 1.75. Although we did not directly compare people who don’t smoke, this finding is broadly consistent with previous studies explicitly demonstrating that people who smoke have a “hyper-resting brain entropy” state (Z. Li et al., 2016).

**Figure 3.**
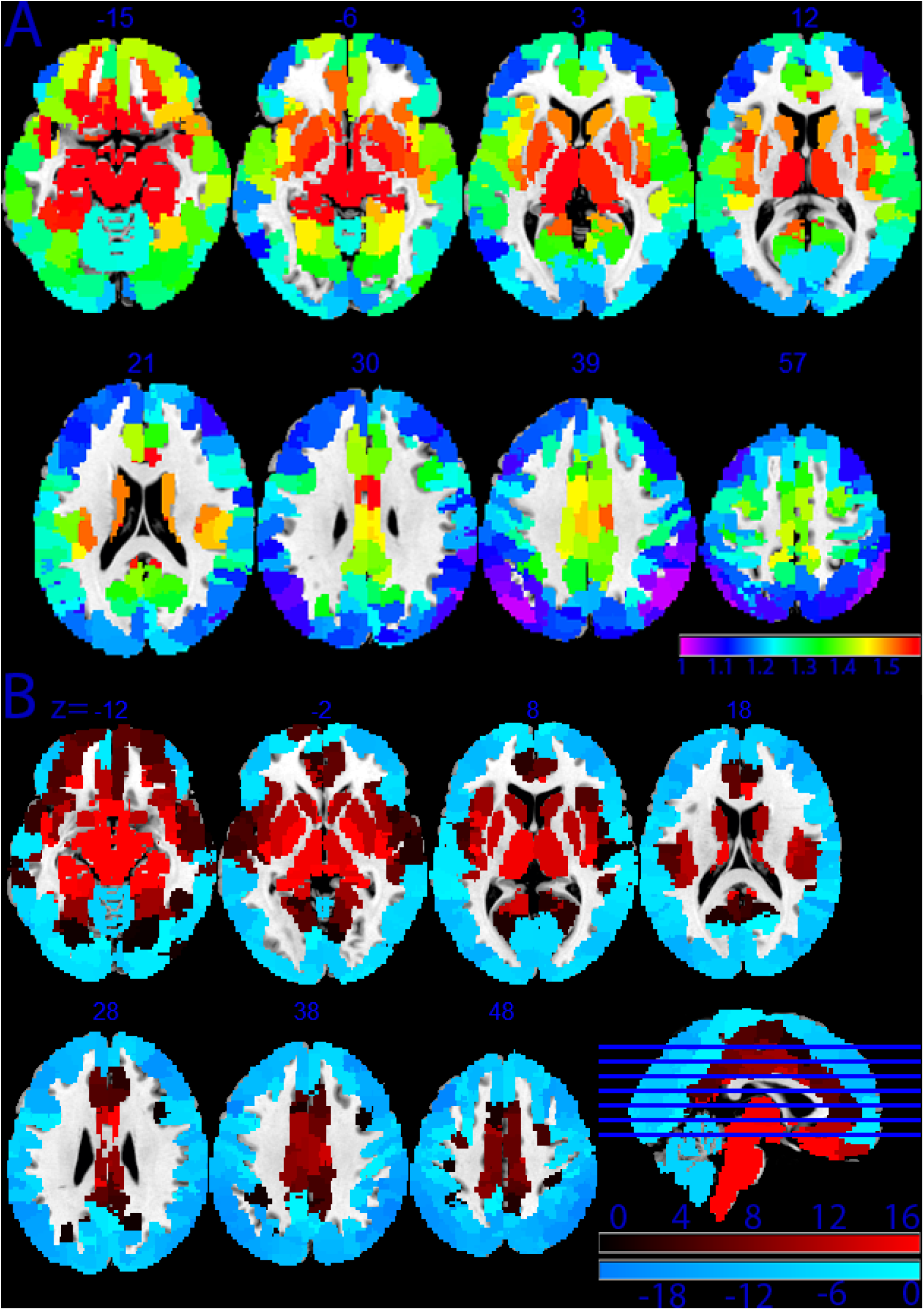
Average Sample Entropy Maps. A) Average sample entropy per region across the brain. Regions are defined by the Schaefer 400 parcellation and 16 sub-cortical regions & 3 cerebellum regions from the Human Connectome Project. Red indicates the highest levels of SampEn observed, and purple indicates the lowest. B) Regions with SampEn above (red) and below (blue) the average SampEn across the brain for people who smoke pre-rTMS stimulation.

## 3.2 Changes in Craving

Participant self-reported craving measurements were collected before and after stimulation to determine if rTMS has an immediate effect on an individual’s cigarette craving. Self-reported measures were compared using a paired-samples t-test and corrected for multiple comparisons using the Bonferroni correction. Craving as measured by the Shiffman-Jarvik Withdrawal Scale craving subscale was found to be significantly different for stimulation to left dlPFC (Pre: 22 (8.25); Post: 20.45 (6.7), t(df) = 2.36(37), *p* = 0.005, Cohen’s *d* = 0.31). No significant differences were found for the Urge to Smoke for stimulation to left dlPFC. No craving measures were found to be significantly different for stimulation to left v5. **Figure 4 & Table 2** show the results for the SJWS-Craving scores for both sessions. Results for the Urge to Smoke are in the supplementary materials.

**Figure 4.**
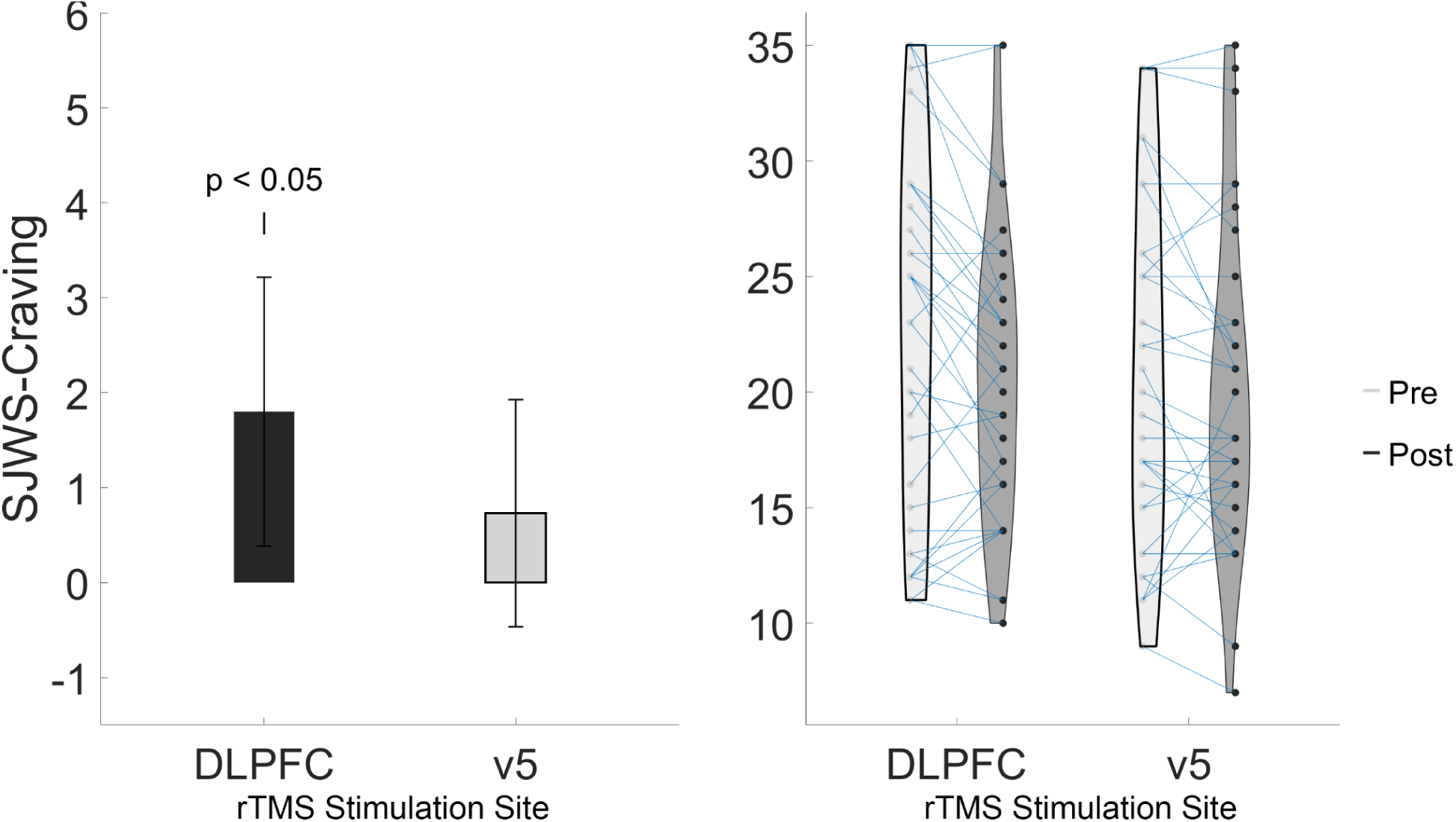
Stimulation to left dlPFC reduced craving in participants. Left) Change score for Shiffman-Jarvik Withdrawal Craving measure from Pre-rTMS minus Post-rTMS values, resulting in larger positive value corresponding to larger reductions in craving. Right) Violin individual point distribution plot, show each participant’s individual change from Pre to Post-rTMS.

**Table 2.** Shiffman-Jarvik Withdrawal Craving Scale Measures. Pre and Post-rTMS to both targets showing both their total craving scores and average craving scores before and after treatment.

| Shiffman-Jarvik Withdrawal Scale Measurements<br>*Mean (SD)* |  |  |  |
| --- | --- | --- | --- |
|  |  | dIPFC | v5 |
| Total Craving | Pre-rTMS | 22 (8.25) | 21.79 (7.74) |
|  | Post-rTMS | 20.45 (6.7) | 21.07 (7.14) |
| Average Craving | Pre-rTMS | 4.57 (1.7) | 4.44 (1.57) |
|  | Post-rTMS | 4.22 (1.4) | 4.26 (1.43) |

### 3.3 Brain Entropy changes in left dlPFC and Insula

Extracted time series using the atlas previously described were normalized and entered into the BENtbx to calculate each node’s sample entropy. Node sample entropy was compared for Pre- and Post-rTMS stimulation for all 419 regions using a paired t-test. All results were corrected for multiple comparisons using the False-Discovery Rate method.

Examining the 19 nodes that include bilateral insula and left dlPFC, we found that 17 out of 19 nodes had significant changes in their SampEn measurements from pre- to post-stimulation to left dlPFC. All nodes showed lower SampEn in post-stimulation scans compared to pre-stimulation. **Table 3** shows the results for each node. No insula or dlPFC nodes were found to change significantly after stimulation to v5. ***Figures 5 & 6*** show the mean SampEn measurements before and after stimulation in the nodes with the most significant change for each stimulation site, and a t-statistic map for those nodes. Figures for all other nodes in these regions can be found in the supplementary materials.

**Figure 5.**
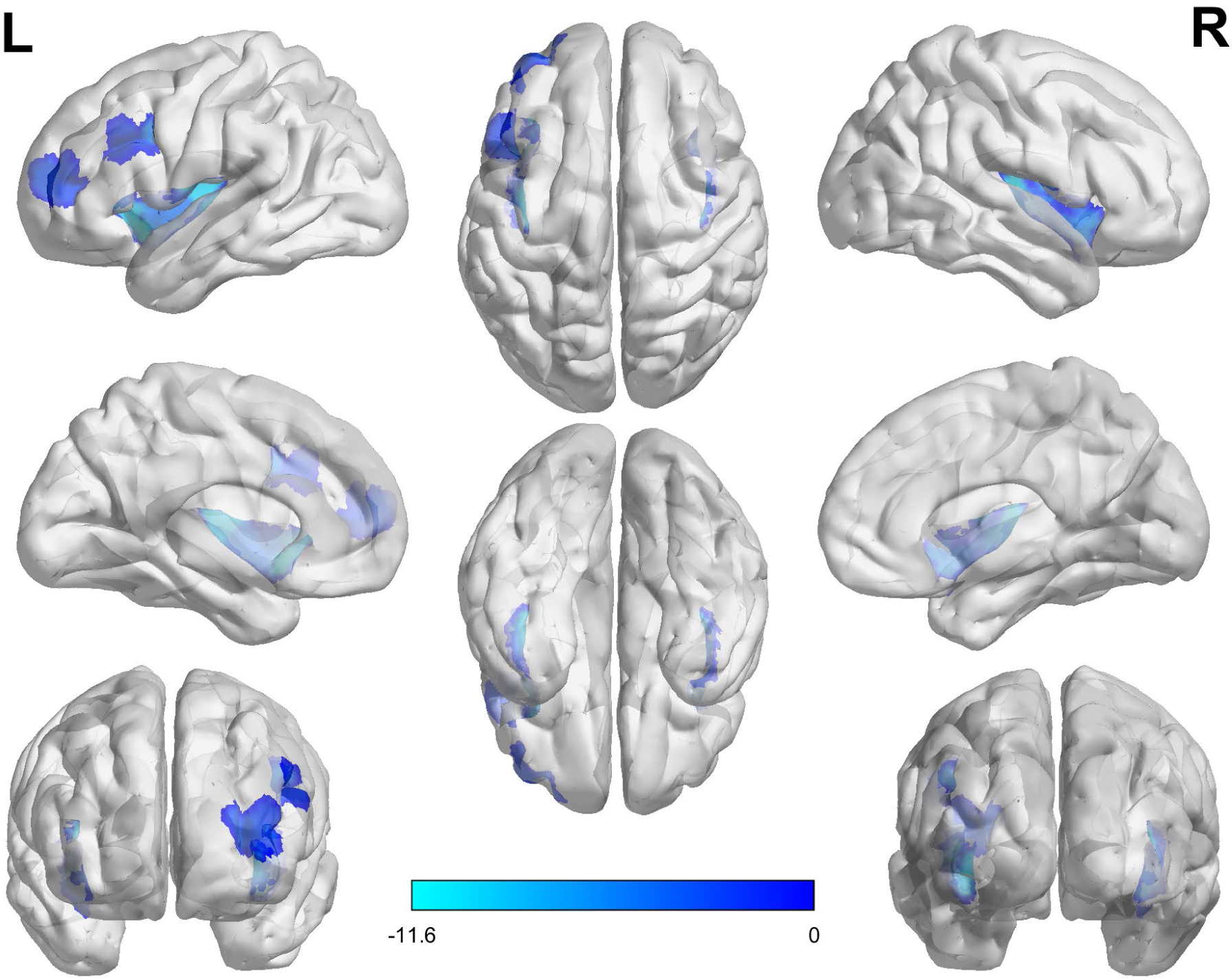
Reductions in entropy in *a priori*-selected ROIs. Regions in blue (insula and left dlPFC) were selected *a priori* as nodes that were expected to show reductions in SampEn as a result of rTMS. The *t*-statistic value for node is shown in blue, with darker values indicating a t-statistic closer to zero, and lighter blues showing increasingly more negative t-statistics as a result of stimulation to left dlPFC, indicating greater rTMS-induced decreases in SampEn.

**Figure 6.**
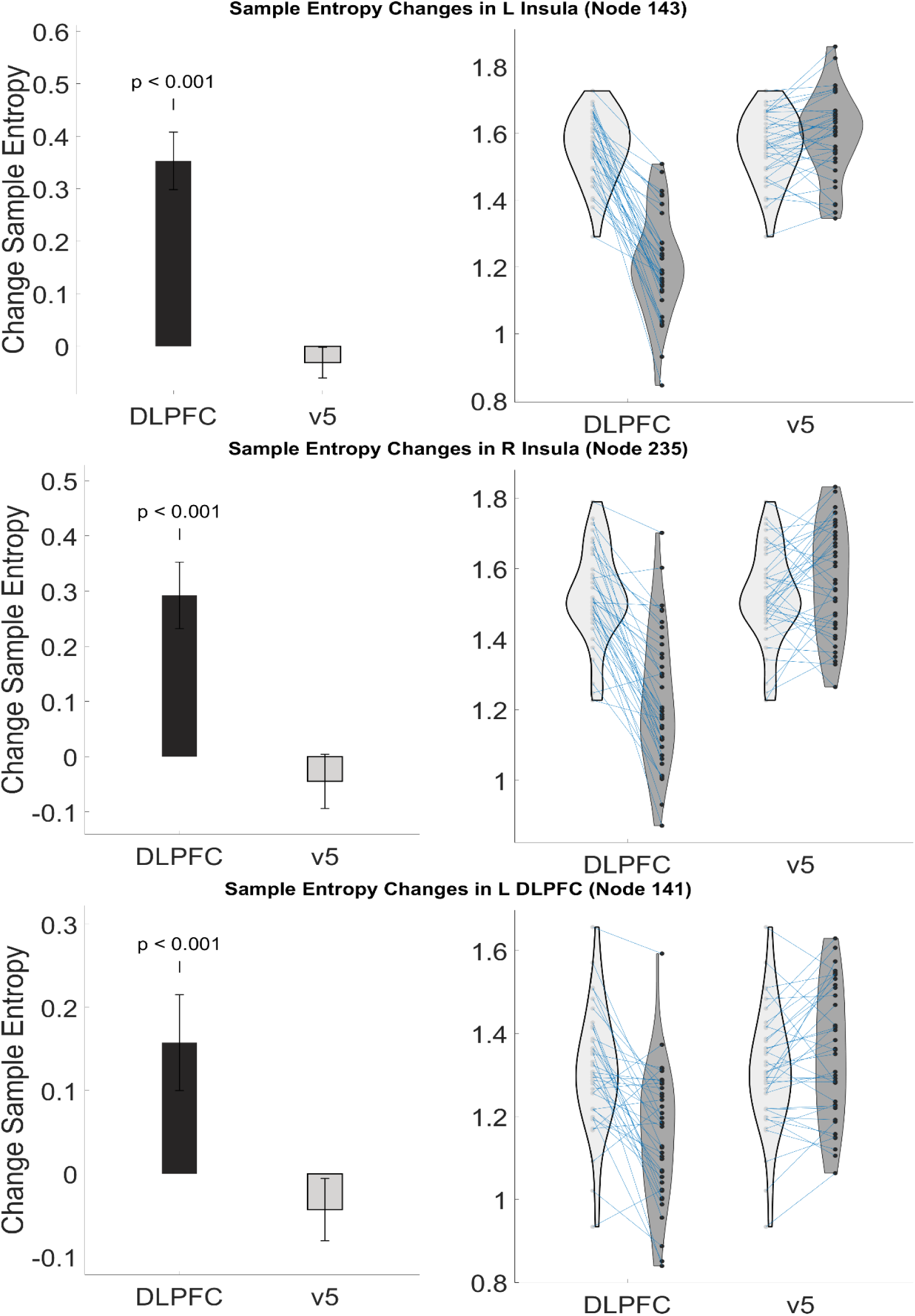
Stimulation to left dlPFC reduced sample entropy in L/R Insula and L dlPFC nodes. Plots show change group in sample entropy (left) and individual changes/distribution (right) for each node that had the lowest p-value and the region that correlated with the node. Change values were calculated by subtracting Post-rTMS entropy values from Pre-rTMS values.

**Table 3.** Sample Entropy T-test Results for Pre vs Post. T-stat, FDR-corrected p-value, and cohen’s d for all a-priori nodes.

| Sample Entropy Statistical Results |  |  |  |
| --- | --- | --- | --- |
| Schaefer Atlas Node | T-Stat (df) | p-val_FDR | Cohen's d |
| 35 (L Insula) | -9.71(41) | 5.06E-11 | 1.82 |
| 98 (L Insula) | -4.76(41) | 4.92E-05 | 0.73 |
| 99 (L Insula) | -9.30(41) | 1.47E-10 | 1.74 |
| 100 (L Insula) | -9.84(41) | 3.92E-11 | 1.84 |
| 143 (L Insula) | -11.51(41) | 1.07E-12 | 3.02 |
| 234 (R Insula) | -10.2(41) | 1.42E-11 | 1.89 |
| 235 (R Insula) | -10.72(41) | 4.49E-12 | 1.83 |
| 236 (R Insula) | -10.39(41) | 9.45E-12 | 1.66 |
| 302 (R Insula) | -6.39(41) | 4.26E-07 | 1.17 |
| 303 (R Insula) | -5.02(41) | 2.32E-05 | 0.97 |
| 304 (R Insula) | -7.28(41) | 3.63E-08 | 1.28 |
| 305 (R Insula) | -8.03(41) | 4.60E-09 | 1.63 |
| 340 (R Insula) | -6.69(41) | 1.86E-07 | 1.22 |
| 137 (L dlPFC) | -2.34(41) | 0.03 | 0.41 |
| 138 (L dlPFC) | -3.06(41) | 0.01 | 0.54 |
| 139 (L dlPFC) | -1.84(41) | 0.09 | 0.44 |
| 140 (L dlPFC) | -2.2(41) | 0.04 | 0.37 |
| 141 (L dlPFC) | -5.85(41) | 2.16E-06 | 1.02 |
| 142 (L dlPFC) | -9.71(41) | 5.06E-11 | 1.82 |

To determine if changes in SampEn influenced the observed changes in an individual’s craving, we correlated each participant’s change in SampEn measurements with their change in craving scores and restricted correlation values to be above r = 0.2. No correlations were found for any of the a priori nodes.

### 3.4 Potential confounding variables

To determine whether other participant characteristics may have influenced SampEn analyses, five variables (sex, ethnicity, age, years of smoking and education) were examined for relationships with SampEn. Pre-rTMS SampEn and change in SampEn were compared between male and female participants using an independent-samples t-test. Pre-rTMS SampEn was compared between all 8 ethnicity categories using a one-way ANOVA. For sex differences, nodes 99, 100, 303, and 304 were found to have differences between male and female participants with females having higher entropy in all nodes. These differences did not survive multiple comparison corrections, but warrant exploration in future studies. No significant differences were found between ethnicity groups. Supplementary Table S4 showing the average SampEn per node for each ethnic group can be found in the supplementary materials. Pearson correlations were calculated for Age/Years of Smoking/Education level vs. Pre-rTMS SampEn to determine if any of the variables influenced the entropy measurement in our participant population. No significant correlations (p > 0.05) were found between age, years of smoking or education level and Pre-rTMS SampEn in left dlPFC, left or right Insula. A table of non-significant correlation values with each variable and the corresponding p-values can be found in the supplementary materials.

### 3.5 Exploratory Findings

After the above *a priori* ROI results were obtained and determined to have no correlation with observed changes in craving, we decided to examine other nodes for significant changes and correlation with behavior. We observed that for left dlPFC stimulation, entropy changed significantly across the majority of the brain. **Figure 7** shows a t-statistic map showing the t-statistic associated with the comparison of each region’s SampEn before and after stimulation. Three nodes (133, 314, and 318) were found to have significant changes in SampEn (Cohen’s d = 0.56,1.06,0.58; p_FDR_ = 0.0016,2 · 10^−4^, 6.1 · 10^−6^, respectively) and have a moderate correlation with craving. In node 133, which overlaps with the inferior temporal gyrus, changes in SampEn correlated with the changes in SJWS-Craving (*r*(df) = 0.36(40)). Nodes 314 and 318, both in the right superior frontal gyrus (SFG), had moderate correlations between their changes in SampEn and changes in UTS (*r*(df)= 0.39(34) and *r*(df) = 0.43(34), respectively). All tables and figures for these results are in the supplementary materials.

**Figure 7.**
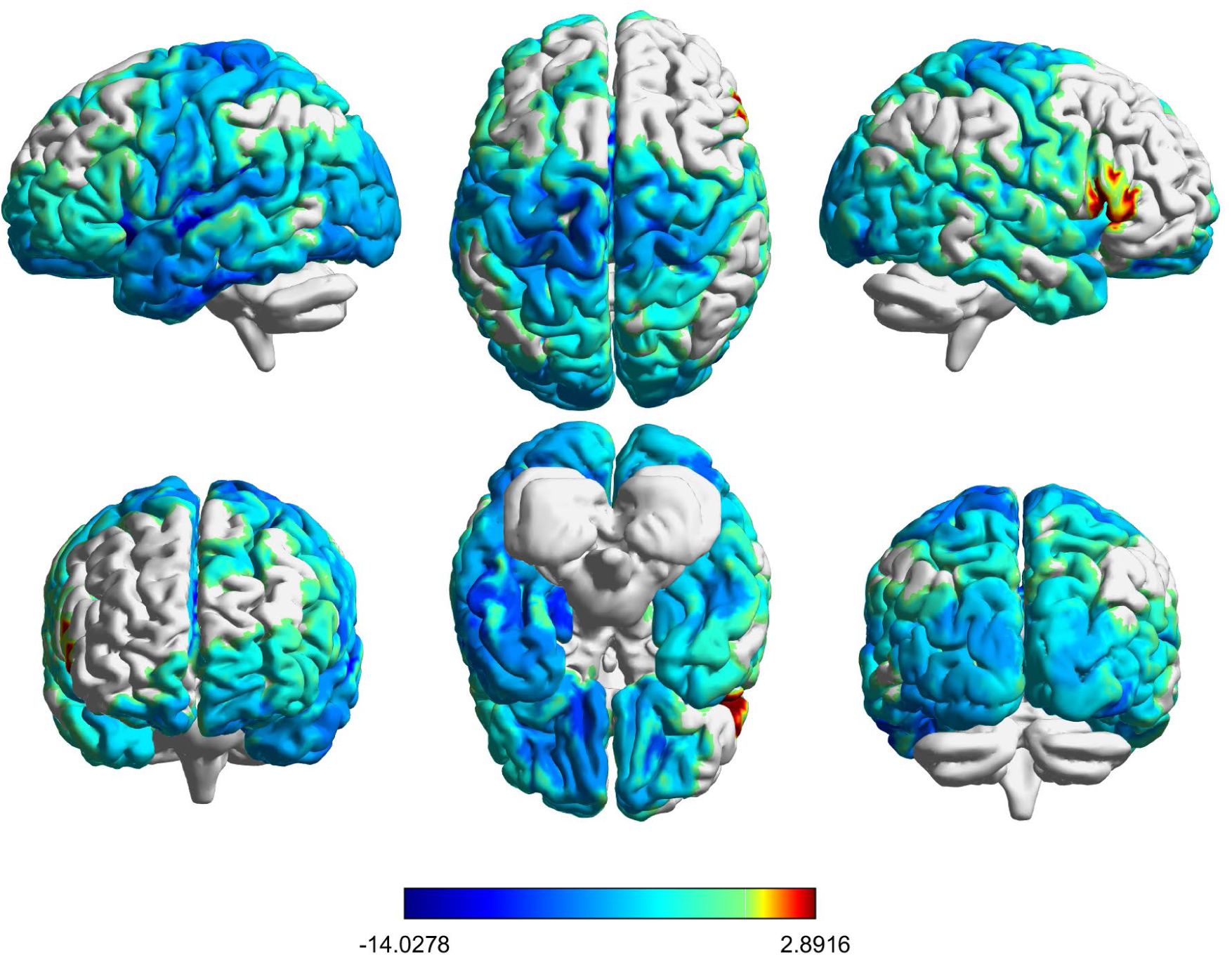
Brainwide changes in SampEn as a result of rTMS to dlPFC show widespread reductions in entropy. Changes in SampEn from pre- to post-dlPFC stimulation are shown here as t-statistics. Red indicates increased entropy following rTMS, gray indicates no change, and increasingly darker blues indicate increasingly greater reductions in entropy as a result of dlPFC stimulation. These reductions in entropy can be observed throughout the brain, with only a few small regions of increased entropy.

## 4. Discussion & Conclusion

### 4.1 dlPFC & Insula

In this study we used excitatory rTMS to the left dlPFC to reduce cigarette craving and sample entropy in the bilateral insula and left dlPFC. Although no correlation was found between the magnitude of SampEn changes in these regions and the magnitude of craving changes, these data suggest that one mechanism by which neuromodulation produces craving relief may include reducing regional brain entropy. This is strengthened by our exploratory results showing that changes in regional SampEn for SFG and ITG did have moderate correlations with craving changes.

This investigation builds on ample previous work strongly relating left dlPFC stimulation to reductions in cigarette craving, consumption, and ultimately, cessation. Although the evidence base for this treatment is mounting, the mechanism by which dlPFC stimulation produces its effects are not well-understood. One small (N = 10) investigation showed that dlPFC stimulation reduces fractional amplitude of low-frequency fluctuations in the insula, and also reduced connectivity between the stimulation site and medial prefrontal cortex (X. Li et al., 2017), suggesting that modulation of insula activity by dlPFC stimulation may be the mechanism by which rTMS alleviates craving.

Our finding that dlPFC stimulation reduces entropy in the insula is consistent with previous evidence suggesting that rTMS to the dlPFC produces its salutary effects. However, based on the current findings, the insula’s link to the observed rTMS induced reductions in craving still remains to be seen. One explanation for this may be that stimulation of the left dlPFC doesn’t cause enough of a reduction in withdrawal and craving to allow for a link to be seen, which may be a result of the relatively small dose of rTMS we delivered in this experiment. Because the magnitude of craving reduction corresponds to the magnitude of entropy reduction in the SFG, even though the difference between pre and post Urge to Smoke scores was minimal and inconsistent in participants, these findings suggest that rTMS to the dlPFC may be a viable target, but not the most effective. Direct stimulation to SFG may prove to be more effective.

The superior frontal gyrus has been linked to smoking through a previous brain stimulation study. (Rose et al., 2011) showed in a small study of 15 participants that excitatory stimulation to SFG resulted in immediate reductions in self-reported craving and reductions in craving due to neutral cues. Two previous studies showed that in people who smoke, SFG demonstrated higher levels of spontaneous activity (Niu et al., 2023) and lower resting functional connectivity (Zhou et al., 2017) relative to controls. These findings could be broadly consistent with our observations of entropy reductions, as rTMS may reduce entropy in SFG, thus causing the region to stabilize its activity and thereby reduce craving.

### 4.2 Limitations

This investigation delivered only single-session rTMS, and therefore conclusions about long-term effects cannot be drawn. Notably, however, in a pivotal multi-center trial of rTMS for smoking cessation, acute (single-session) reductions in craving did predict successful smoking cessation (Zangen et al., 2021). Additionally, our control condition in this investigation involved stimulation to a different brain site that was delivered to our test population of people with TUD. The lack of a control group (people who do not smoke) prevents any conclusions about entropy in people with and without TUD from being drawn from this data; however, previous work has performed this comparison and found higher brain entropy in people who smoke compared to controls (Z. Li et al., 2016).

Brain entropy was calculated in this study using fMRI collected data, which has a lower temporal resolution than other methods of neuroimaging, such as electroencephalography. This restriction on temporal resolution potentially limits the results determined here and should be validated using a neuroimaging method with higher resolution so that more accurate measures of entropy can be determined due to finer time scales. Likewise, although extensive denoising of data was carried out, residual noise could remain in the data and therefore influence the results.

### 4.3 Conclusion

In this study, we were able to replicate previous findings that rTMS can reduce sample entropy in the brain, and extended these findings in people who smoke, showing that the effect of rTMS on sample entropy is consistent across different populations. We also replicated previous observations about the distribution of brain entropy across the brain and observed evidence of potentially increased resting entropy in people who smoke. Although changes to insula and left DLPFC SampEn did not correlate with changes in behavior, we did find that post-TMS reductions in entropy in two other regions, the SFG and ITG, correlated with rTMS-induced reductions in craving. This result provides additional (although indirect) evidence that entropy is higher in people who smoke than people who do not, and suggests that by reducing entropy in specific regions associated with smoking, we can reduce cigarette craving. This work shows that sample entropy may be a potential biomarker for measuring efficacy of rTMS-based smoking cessation treatments.

### 4.4 Future Directions

Future studies should examine this effect in larger populations using more substantial doses of rTMS. Moreover, future investigations may test the effect of using baseline entropy in regions associated with smoking, specifically insula and SFG, to adjust individual treatments. Next, we will also need to explore the functional connectivity changes in these participants to see if the regions found with significant changes in SampEn also have changes in functional connectivity and compare them separately and together as predictors of behavior changes. Expanding upon this work, further investigations into brain complexity should be examined outside of just regional complexity. These should include measures of complexity of functional connections using functional entropy (Yao et al., 2013), entropy states and directional influences of entropy (Varley et al., 2023), and community mapping entropy (Betzel et al., 2019). By developing our understanding of how these measures of entropy change due to TMS, entropy can be better applied as a biomarker for treatments.

## Data and Code availability

The data and code that support the results of this study are available on Github (https://github.com/humanbrainzappingatucla). Any additional information required to reanalyze the data used in this paper is available upon request and use agreement with the corresponding author

## Author Contributions

Conceptualization, T.J. and N.P.; Methodology, T.J. and J.N.; Software, T.J.; Formal Analysis, T.J.; Investigation, M.A., T.J., and N.P.; Data Curation, T.J.; Writing - Original Draft, T.J. and N.P.; Writing - Review & Editing, T.J. M.A. J.N., and N.P.; Visualization, T.J.; Supervision, T.J. and N.P.; Project Administration, M.A. and N.P.; Funding Acquisition, N.P.

## Acknowledgements

We thank the staff of the Center for Cognitive Neuroscience for providing aid and support for all fMRI imaging sessions. We thank the UCLA TMS Clinical and Research Services for providing technical and medical support during all stimulation sessions. We are grateful to Lucina Uddin for helpful discussions and feedback. We thank Anthony Sun, Melanie Beltran and Riley Russell for assisting the investigators.

## Supplemental Materials

Supplemental figures and tables can be found here: (link to be generated)

## Supplemental Materials

### Urge to Smoke Results

**Figure S1.**
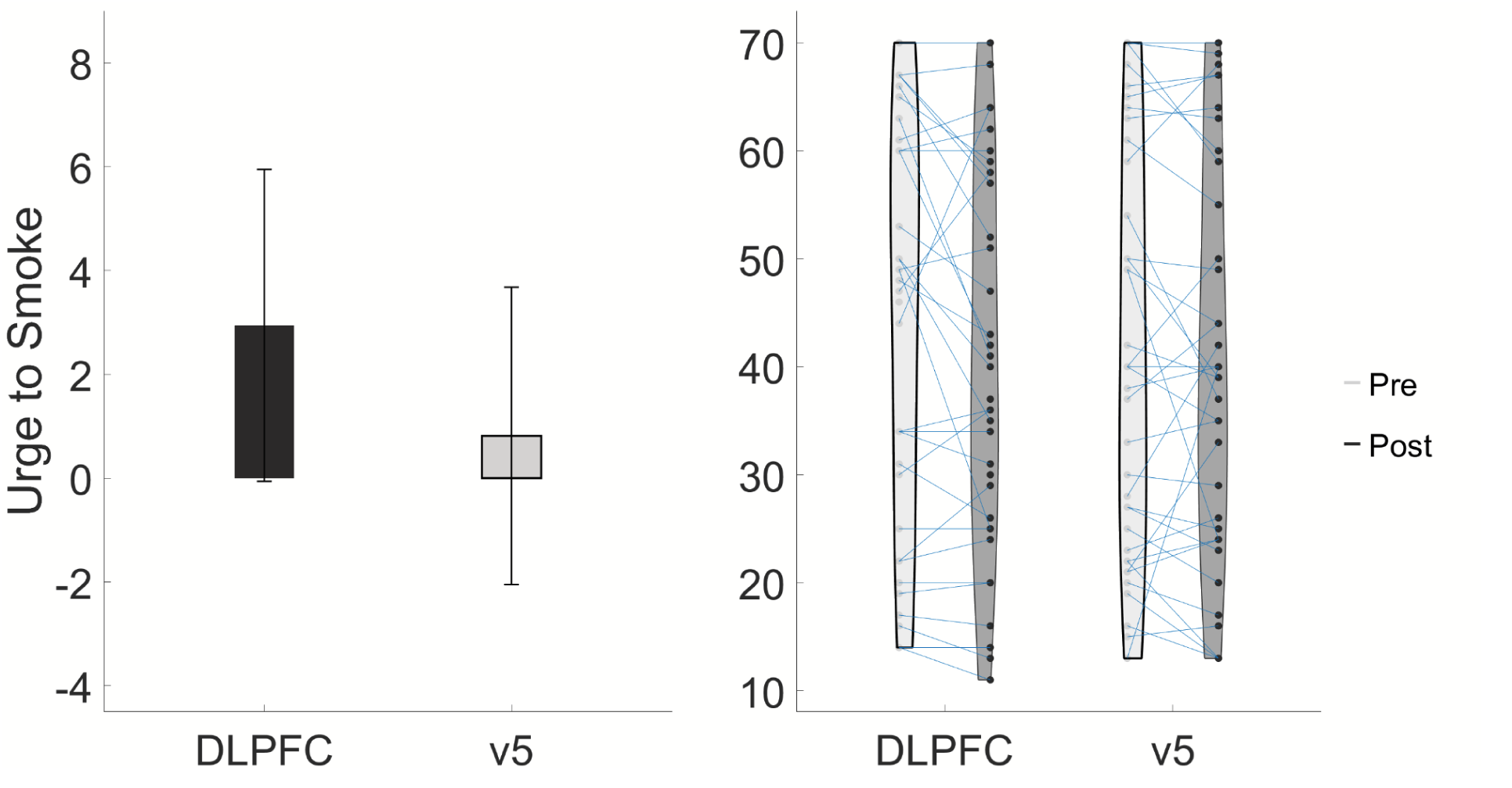
Changes in Urge to Smoke (related to Figure 3) No significant changes in Urge to Smoking were found for stimulation to left dlPFC.

**Table S1.** Urge to Smoke measurements (related to Table 1) Pre and Post-rTMS to both targets showing both their urge to smoke scores before and after treatment.

| Urge to Smoke Measurements |  |  |  |
| --- | --- | --- | --- |
|  |  | dIPFC | v5 |
| UTS | Pre-rTMS | 43.74 (20.05) | 43.4 (19.54) |
|  | Post-rTMS | 41.1 (19.1) | 41.7 (18.26) |

### Sample Entropy of L/R Insula and left dlPFC nodes

**Table S2.** Sample Entropy of each node Pre and Post-rTMS (related to Figure 5) Sample entropy measures for each node found to have significant changes in sample entropy post-rTMS to DLPFC. All measures are given as mean measures with standard deviation.

| Sample Entropy Pre-rTMS & Post-rTMS |  |  |  |
| --- | --- | --- | --- |
| Schaefer Atlas Node | Time | dIPFC entropy, mean (SD) | V5 entropy, mean (SD) |
| 35 (L Insula) | Pre-rTMS | 1.516 (0.159) | 1.528 (0.156) |
|  | Post-rTMS | 1.191 (0.193) | 1.539 (0.17) |
| 98 (L Insula) | Pre-rTMS | 1.463 (0.114) | 1.466 (0.123) |
|  | Post-rTMS | 1.324 (0.199) | 1.442 (0.147) |
| 99 (L Insula) | Pre-rTMS | 1.464 (0.128) | 1.481 (0.109) |
|  | Post-rTMS | 1.233 (0.162) | 1.493 (0.143) |
| 100 (L Insula) | Pre-rTMS | 1.522 (0.119) | 1.529 (0.111) |
|  | Post-rTMS | 1.246 (0.182) | 1.567 (0.127) |
| 143 (L Insula) | Pre-rTMS | 1.533 (0.134) | 1.553 (0.111) |
|  | Post-rTMS | 1.208 (0.141) | 1.586 (0.139) |
| 234 (R Insula) | Pre-rTMS | 1.563 (0.135) | 1.578 (0.12) |
|  | Post-rTMS | 1.26 (0.184) | 1.586 (0.137) |
| 235 (R Insula) | Pre-rTMS | 1.515 (0.13) | 1.528 (0.137) |
|  | Post-rTMS | 1.22 (0.186) | 1.563 (0.151) |
| 236 (R Insula) | Pre-rTMS | 1.546 (0.14) | 1.564 (0.14) |
|  | Post-rTMS | 1.245 (0.197) | 1.596 (0.208) |
| 302 (R Insula) | Pre-rTMS | 1.596 (0.112) | 1.59 (0.119) |
|  | Post-rTMS | 1.429 (0.182) | 1.617 (0.131) |
| 303 (R Insula) | Pre-rTMS | 1.467 (0.113) | 1.484 (0.112) |
|  | Post-rTMS | 1.347 (0.155) | 1.484 (0.121) |
| 304 (R Insula) | Pre-rTMS | 1.537 (0.095) | 1.544 (0.095) |
|  | Post-rTMS | 1.357 (0.158) | 1.552 (0.13) |
| 305 (R Insula) | Pre-rTMS | 1.517 (0.129) | 1.522 (0.129) |
|  | Post-rTMS | 1.308 (0.166) | 1.567 (0.123) |
| 340 (R Insula) | Pre-rTMS | 1.527 (0.137) | 1.525 (0.128) |
|  | Post-rTMS | 1.379 (0.144) | 1.548 (0.141) |
| 137 (L dlPFC) | Pre-rTMS | 1.138 (0.129) | 1.149 (0.114) |
|  | Post-rTMS | 1.083 (0.138) | 1.13 (0.147) |
| 138 (L dlPFC) | Pre-rTMS | 1.167 (0.142) | 1.17 (0.131) |
|  | Post-rTMS | 1.097 (0.147) | 1.15 (0.134) |
| 139 (L dlPFC) | Pre-rTMS | 1.121 (0.105) | 1.127 (0.1) |
|  | Post-rTMS | 1.088 (0.137) | 1.119 (0.127) |
| 140 (L dlPFC) | Pre-rTMS | 1.142 (0.134) | 1.158 (0.124) |
|  | Post-rTMS | 1.094 (0.134) | 1.16 (0.119) |
| 141 (L dlPFC) | Pre-rTMS | 1.306 (0.146) | 1.308 (0.14) |
|  | Post-rTMS | 1.152 (0.151) | 1.348 (0.155) |
| 142 (L dlPFC) | Pre-rTMS | 1.15 (0.153) | 1.153 (0.141) |
|  | Post-rTMS | 1.11 (0.135) | 1.155 (0.124) |

**Figure S2.**
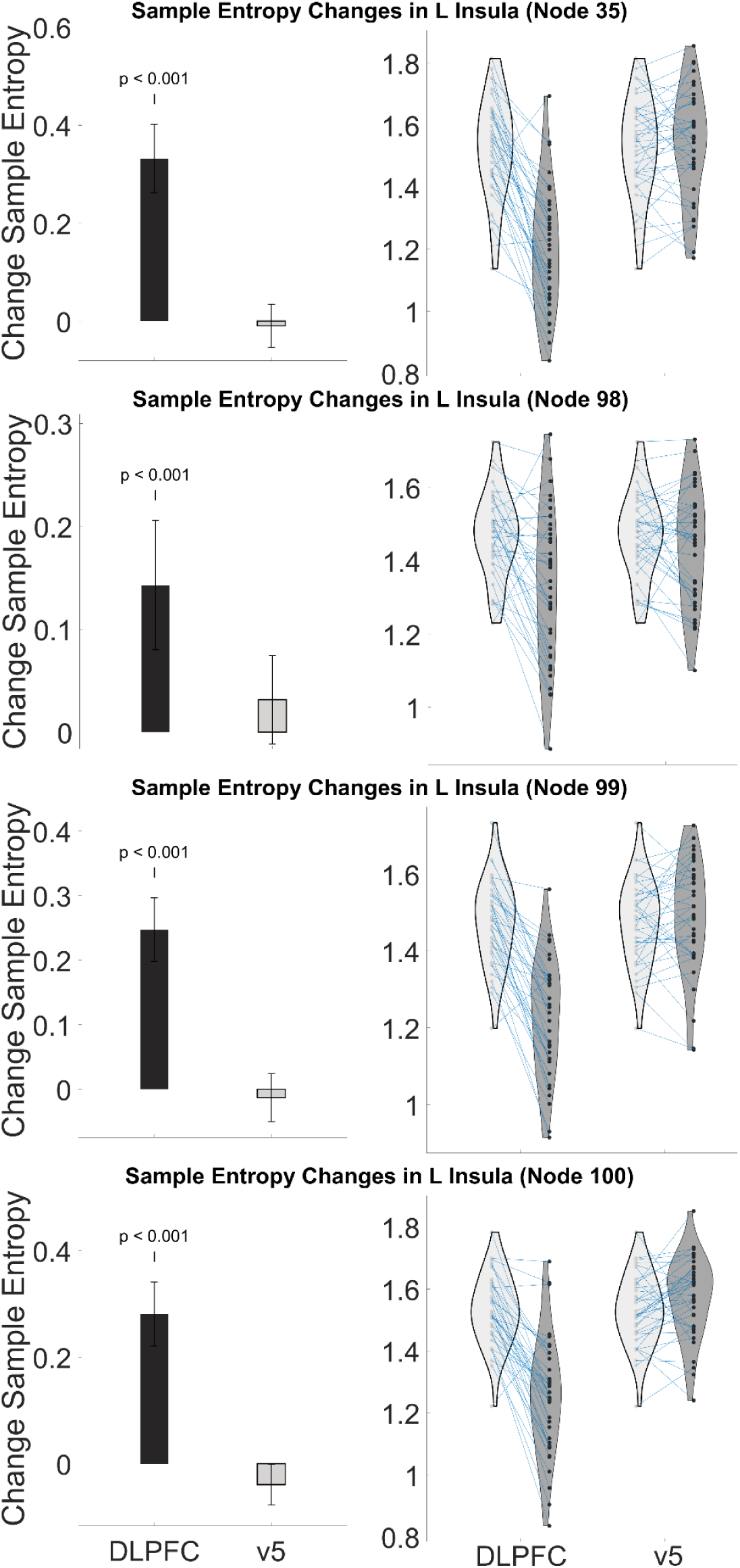
Stimulation to left dlPFC reduced sample entropy in left Insula (related to Figure 5) Plots show change group in sample entropy (left) and individual changes/distribution (right) for each left Insula node. Change values were calculated by subtracting Post-rTMS entropy values from Pre-rTMS values.

**Figure S3.**
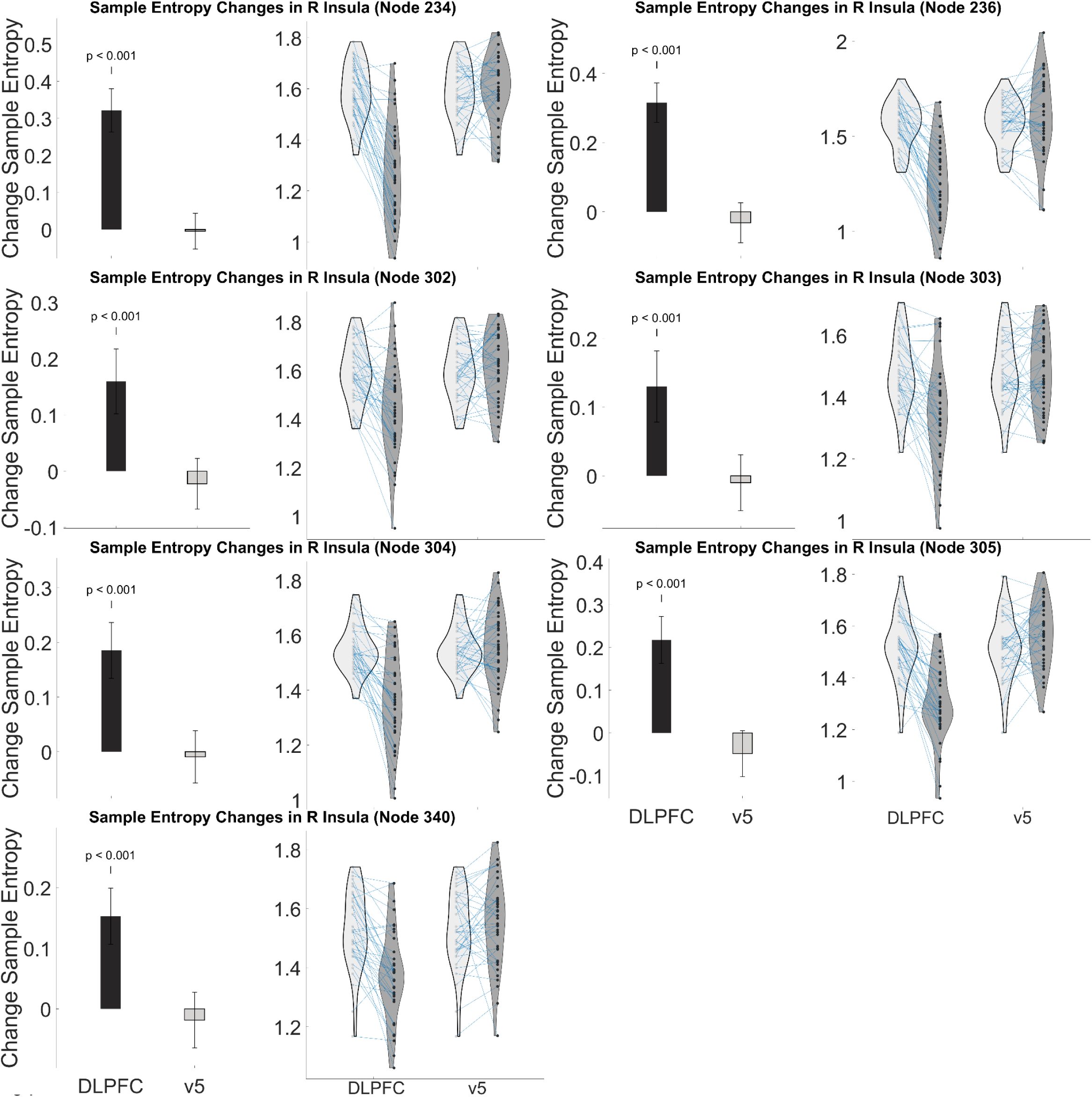
Stimulation to left dlPFC reduced sample entropy in right Insula (related to Figure 5) Plots show change group in sample entropy (left) and individual changes/distribution (right) for each right Insula node. Change values were calculated by subtracting Post-rTMS entropy values from Pre-rTMS values.

**Figure S4.**
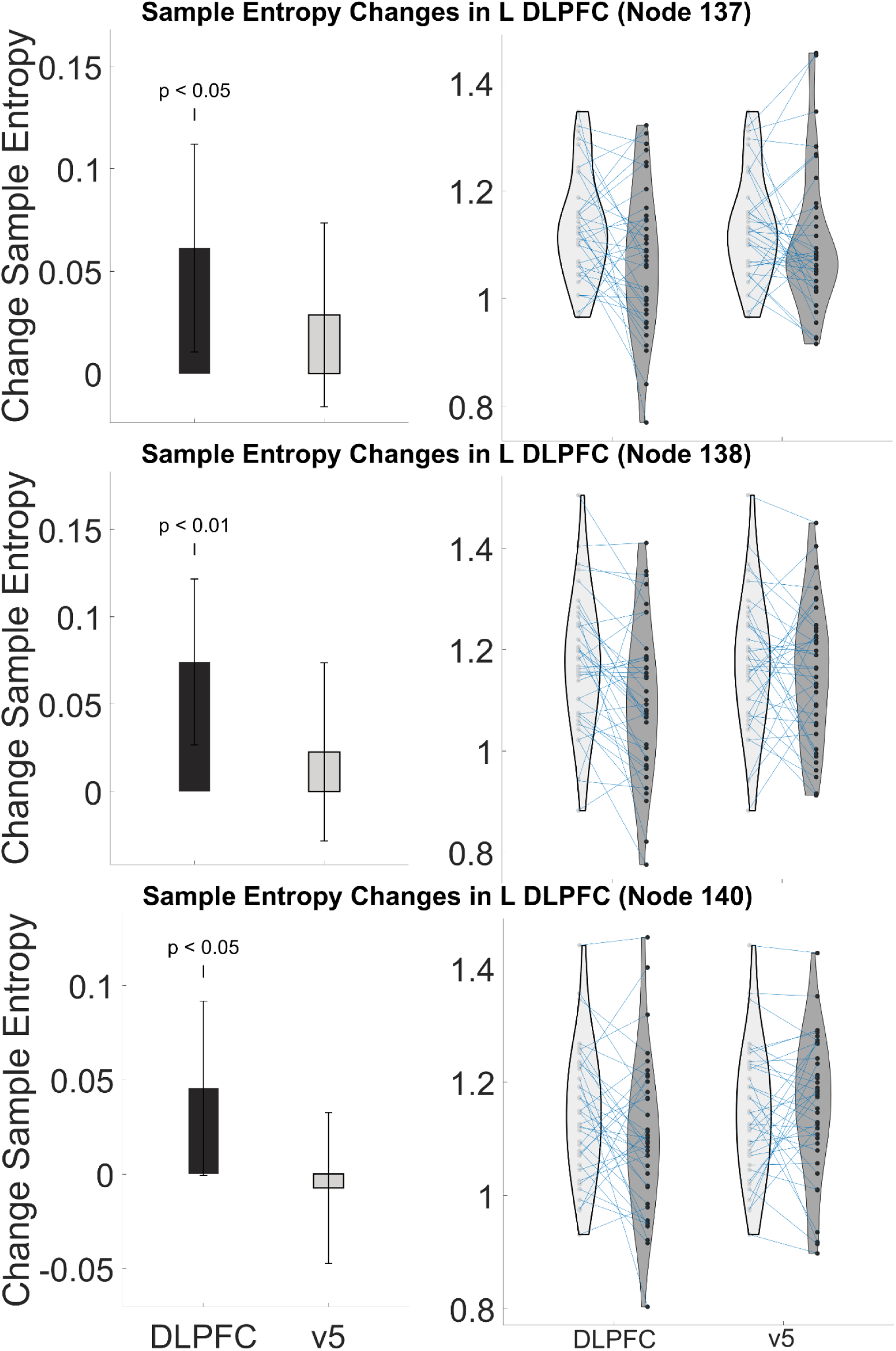
Stimulation to left dlPFC reduced sample entropy in left Insula (related to Figure 5) Plots show change group in sample entropy (left) and individual changes/distribution (right) for each left dlPFC node. Change values were calculated by subtracting Post-rTMS entropy values from Pre-rTMS values.

### Confounding Variable Correlations

**Table S3.** No significant differences in Pre-rTMS entropy between ethnicities. Ethnic groups were compared for Pre-rTMS sample entropy measures to determine if there were significant differences. No significant differences were found for any of the nodes. This Table Shows how many participants in each ethnic group were included in this study and their group’s mean sample entropy with standard deviation for each node.

| Sample Entropy across Ethnicity |  |  |  |  |  |  |
| --- | --- | --- | --- | --- | --- | --- |
| Node | Asian<br>(N=5) | Native<br>Hawaiian/<br>Pacific<br>Islander<br>(N=1) | Black /<br>African<br>American<br>(N=11) | White<br>(N=19) | Hispanic<br>(N=4) | More than<br>One Race<br>(N=2) |
| 35 | 1.6(0.2) | 1.4(0) | 1.5(0.2) | 1.5(0.2) | 1.5(0.1) | 1.5(0.1) |
| 98 | 1.5(0.1) | 1.4(0) | 1.5(0.1) | 1.4(0.1) | 1.5(0.1) | 1.5(0.1) |
| 99 | 1.5(0.1) | 1.5(0) | 1.5(0.1) | 1.4(0.1) | 1.6(0.1) | 1.4(0) |
| 100 | 1.6(0.1) | 1.5(0) | 1.5(0.1) | 1.5(0.1) | 1.5(0.1) | 1.5(0) |
| 143 | 1.6(0.1) | 1.6(0) | 1.5(0.2) | 1.5(0.1) | 1.5(0.1) | 1.6(0.1) |
| 234 | 1.6(0.2) | 1.5(0) | 1.5(0.2) | 1.5(0.1) | 1.6(0.1) | 1.6(0) |
| 235 | 1.5(0) | 1.4(0) | 1.5(0.2) | 1.5(0.1) | 1.5(0) | 1.5(0) |
| 236 | 1.6(0.1) | 1.6(0) | 1.5(0.1) | 1.5(0.2) | 1.6(0.1) | 1.5(0.1) |
| 302 | 1.7(0.1) | 1.4(0) | 1.6(0.1) | 1.6(0.1) | 1.6(0.1) | 1.5(0) |
| 303 | 1.5(0.1) | 1.4(0) | 1.5(0.1) | 1.4(0.1) | 1.5(0.1) | 1.5(0.2) |
| 304 | 1.6(0.1) | 1.5(0) | 1.5(0.1) | 1.5(0.1) | 1.6(0.1) | 1.5(0) |
| 305 | 1.6(0.1) | 1.4(0) | 1.5(0.2) | 1.5(0.1) | 1.5(0) | 1.5(0) |
| 340 | 1.6(0.1) | 1.5(0) | 1.5(0.1) | 1.5(0.1) | 1.6(0.1) | 1.6(0.1) |
| 137 | 1.2(0.2) | 1.1(0) | 1.1(0.2) | 1.1(0.1) | 1.1(0) | 1.1(0.1) |
| 138 | 1.3(0.1) | 1.3(0) | 1.1(0.2) | 1.1(0.1) | 1.2(0.1) | 1.2(0) |
| 139 | 1.1(0.1) | 1.1(0) | 1.1(0.1) | 1.1(0.1) | 1.1(0) | 1.1(0) |
| 140 | 1.2(0.1) | 1.1(0) | 1.2(0.2) | 1.1(0.1) | 1.2(0.1) | 1.1(0.1) |
| 141 | 1.4(0.1) | 1.4(0) | 1.3(0.2) | 1.2(0.1) | 1.4(0.1) | 1.3(0.1) |
| 142 | 1.3(0.1) | 1.2(0) | 1.1(0.2) | 1.1(0.1) | 1.3(0.1) | 1.1(0.1) |

**Table S4.**
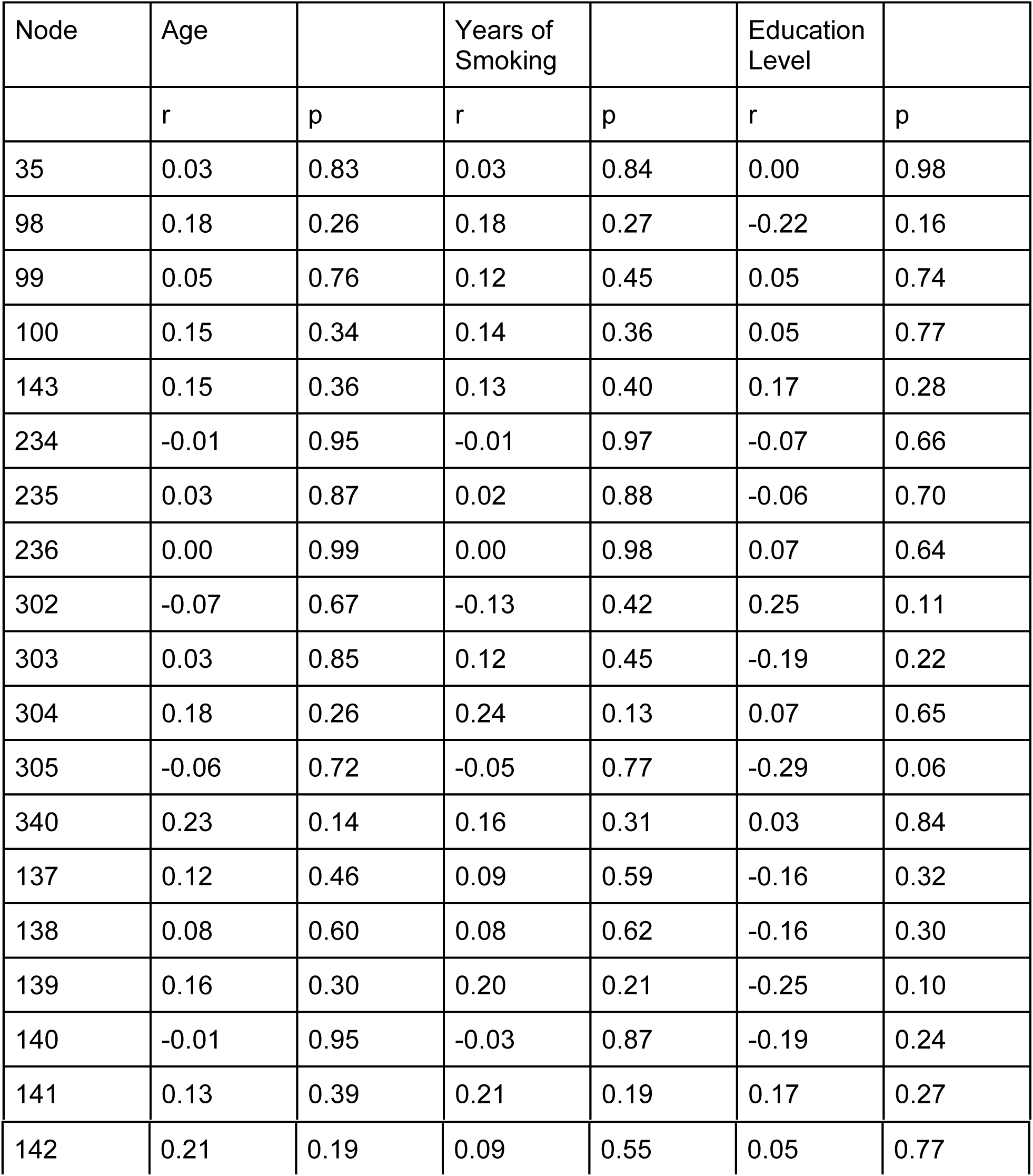
No significant correlations between Pre-rTMS entropy and confounding variables. Pearson correlations for three confounding variables (age, years of smoking, and education) were calculation for Pre-rTMS sample entropy measures to determine if there were significant correlations. No correlations were found for any of the nodes for any of the variables. This Table Shows the pearson correlation coefficient (r) and the p-value of each coefficient for each variable with pre-rTMS sample entropy measures.

| Node | Age |  | Years of Smoking |  | Education Level |  |
| --- | --- | --- | --- | --- | --- | --- |
|  | r | p | r | p | r | p |
| 35 | 0.03 | 0.83 | 0.03 | 0.84 | 0.00 | 0.98 |
| 98 | 0.18 | 0.26 | 0.18 | 0.27 | -0.22 | 0.16 |
| 99 | 0.05 | 0.76 | 0.12 | 0.45 | 0.05 | 0.74 |
| 100 | 0.15 | 0.34 | 0.14 | 0.36 | 0.05 | 0.77 |
| 143 | 0.15 | 0.36 | 0.13 | 0.40 | 0.17 | 0.28 |
| 234 | -0.01 | 0.95 | -0.01 | 0.97 | -0.07 | 0.66 |
| 235 | 0.03 | 0.87 | 0.02 | 0.88 | -0.06 | 0.70 |
| 236 | 0.00 | 0.99 | 0.00 | 0.98 | 0.07 | 0.64 |
| 302 | -0.07 | 0.67 | -0.13 | 0.42 | 0.25 | 0.11 |
| 303 | 0.03 | 0.85 | 0.12 | 0.45 | -0.19 | 0.22 |
| 304 | 0.18 | 0.26 | 0.24 | 0.13 | 0.07 | 0.65 |
| 305 | -0.06 | 0.72 | -0.05 | 0.77 | -0.29 | 0.06 |
| 340 | 0.23 | 0.14 | 0.16 | 0.31 | 0.03 | 0.84 |
| 137 | 0.12 | 0.46 | 0.09 | 0.59 | -0.16 | 0.32 |
| 138 | 0.08 | 0.60 | 0.08 | 0.62 | -0.16 | 0.30 |
| 139 | 0.16 | 0.30 | 0.20 | 0.21 | -0.25 | 0.10 |
| 140 | -0.01 | 0.95 | -0.03 | 0.87 | -0.19 | 0.24 |
| 141 | 0.13 | 0.39 | 0.21 | 0.19 | 0.17 | 0.27 |
| 142 | 0.21 | 0.19 | 0.09 | 0.55 | 0.05 | 0.77 |

### Exploratory Findings Entropy Results

**Table S5.**
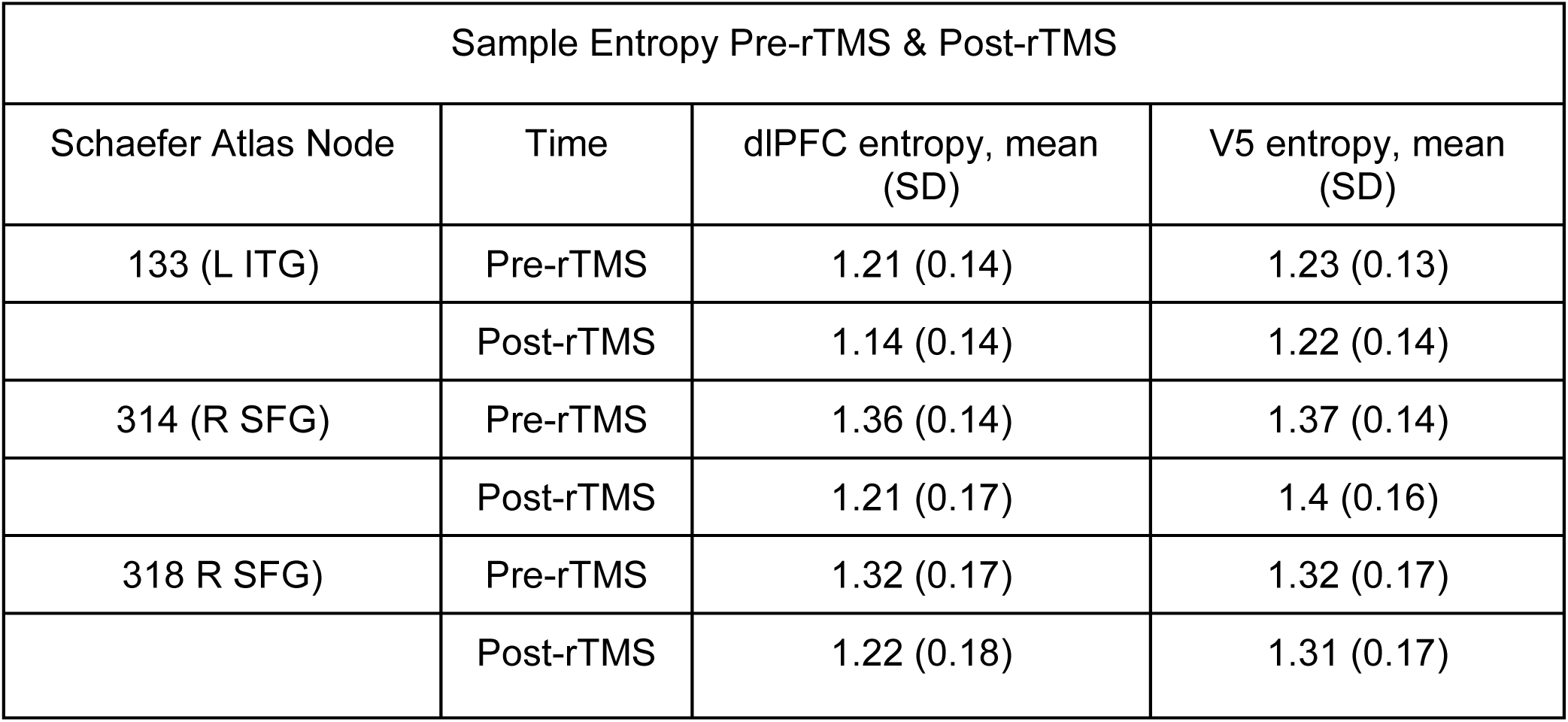
Sample Entropy of each exploratory node Pre and Post-rTMS (related to Figure 7) Sample entropy measures for each node found to have significant changes in sample entropy post-rTMS to DLPFC. All measures are given as mean measures with standard deviation.

| Sample Entropy Pre-rTMS & Post-rTMS |  |  |  |
| --- | --- | --- | --- |
| Schaefer Atlas Node | Time | dIPFC entropy, mean (SD) | V5 entropy, mean (SD) |
| 133 (L ITG) | Pre-rTMS | 1.21 (0.14) | 1.23 (0.13) |
|  | Post-rTMS | 1.14 (0.14) | 1.22 (0.14) |
| 314 (R SFG) | Pre-rTMS | 1.36 (0.14) | 1.37 (0.14) |
|  | Post-rTMS | 1.21 (0.17) | 1.4 (0.16) |
| 318 R SFG) | Pre-rTMS | 1.32 (0.17) | 1.32 (0.17) |
|  | Post-rTMS | 1.22 (0.18) | 1.31 (0.17) |

**Figure S5.**
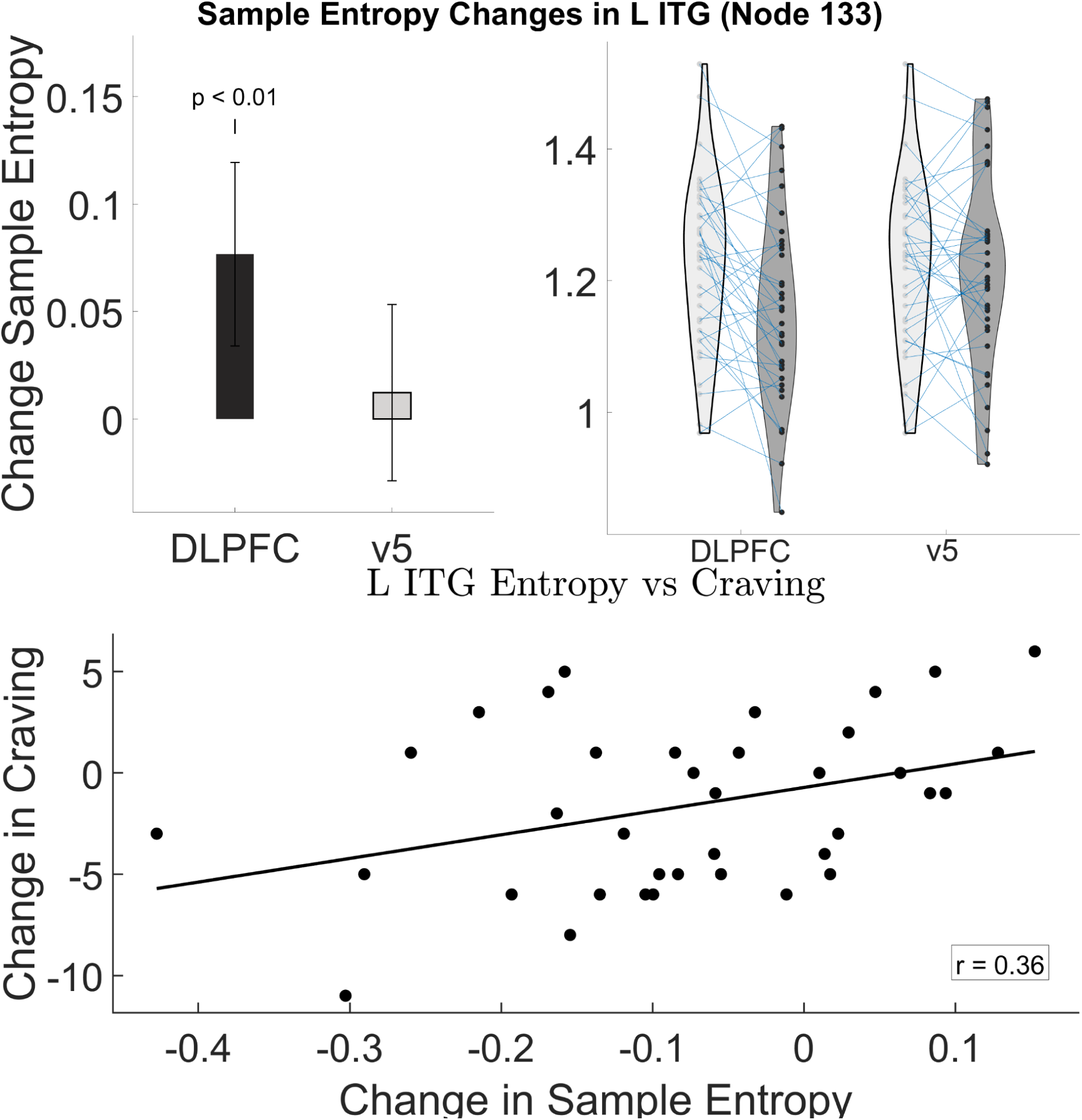
Stimulation to left dlPFC reduced sample entropy in left Inferior Temporal Gyrus (ITG) (related to Figure 7) Top: Plots show change group in sample entropy (left) and individual changes/distribution (right) for this node and the region that correlated with the node. Change values were calculated by subtracting Post-rTMS entropy values from Pre-rTMS values. **Bottom:** Correlation plot between changes in sample entropy in left ITG and craving (r=0.36, p= 0.027)as measured by the Shiffman-Jarvik Withdrawal Scale (SJWS) subscale for craving.

**Figure S6.**
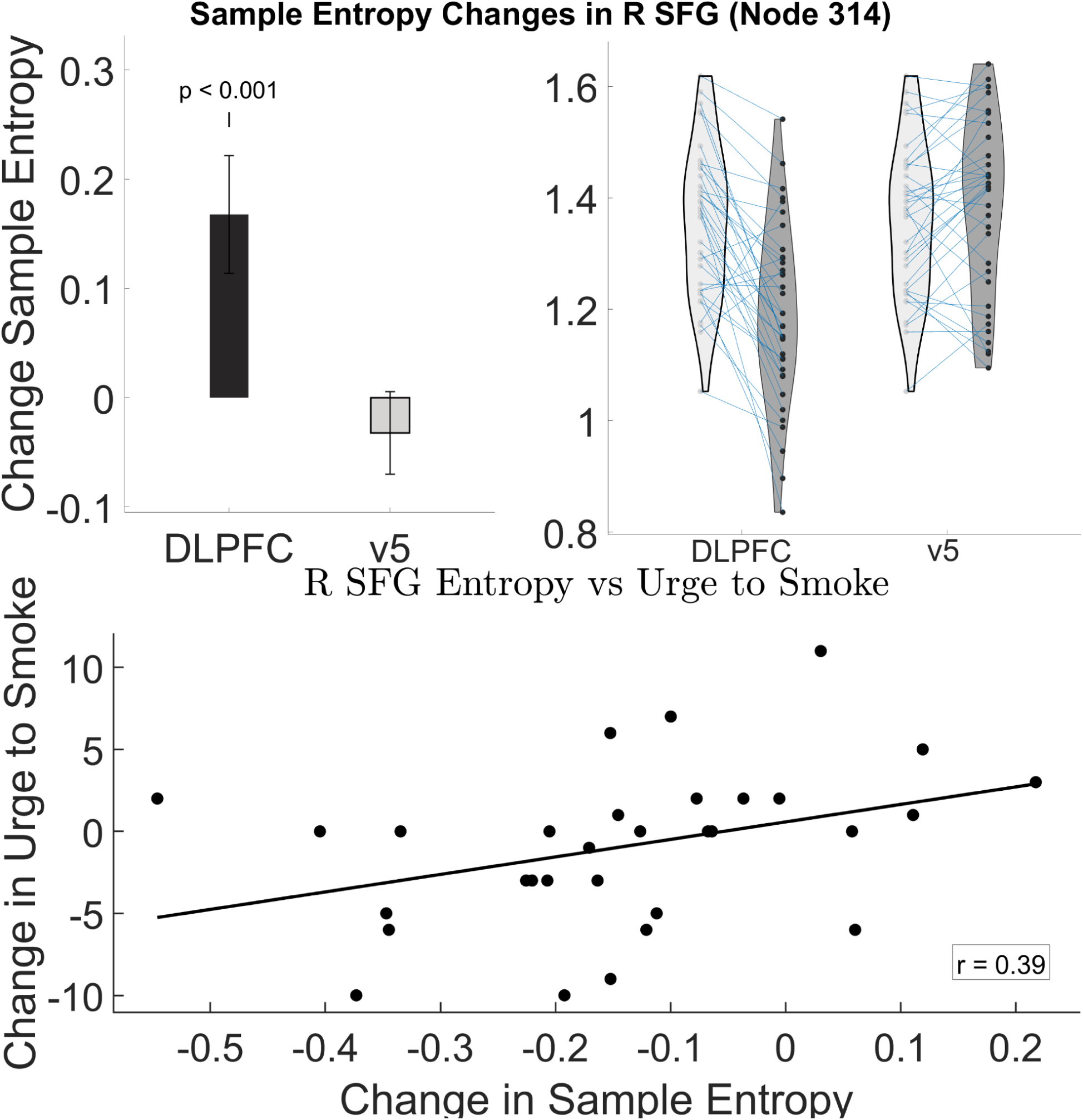
Stimulation to left dlPFC reduced sample entropy in right superior frontal gyrus (SFG)Top: Plots show change group in sample entropy (left) and individual changes/distribution (right) for this node and the region that correlated with the node. Change values were calculated by subtracting Post-rTMS entropy values from Pre-rTMS values. **Bottom:** Correlation plot between changes in sample entropy in right SFG and craving (r=0.39, p= 0.025)as measured by the Shiffman-Jarvik Withdrawal Scale (SJWS) subscale for craving.

**Figure S7.**
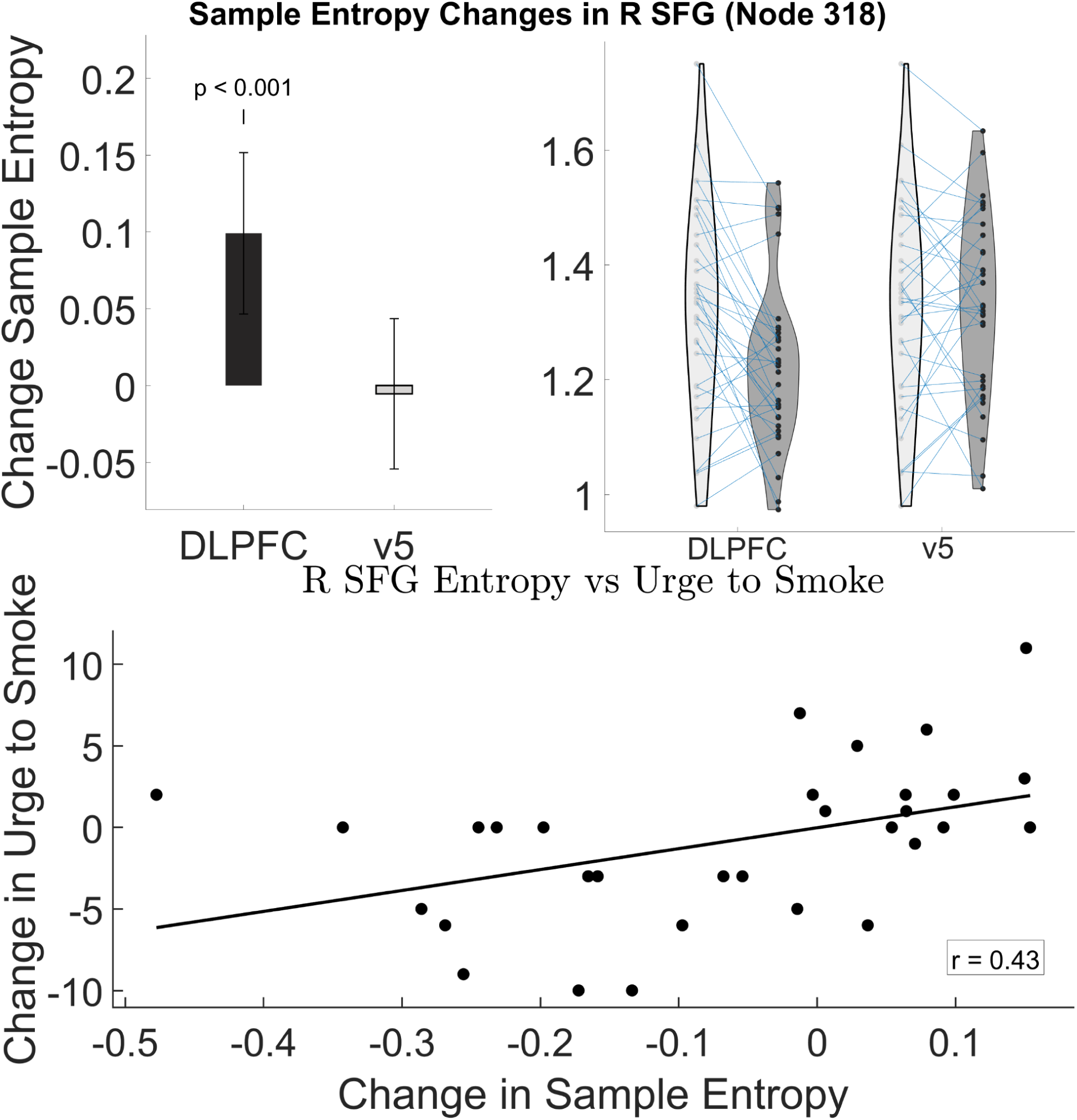
Stimulation to left dlPFC reduced sample entropy in right superior frontal gyrus (SFG) Top: Plots show change group in sample entropy (left) and individual changes/distribution (right) for this node and the region that correlated with the node. Change values were calculated by subtracting Post-rTMS entropy values from Pre-rTMS values. **Bottom:** Correlation plot between changes in sample entropy in right SFG and craving (r=0.43, p= 0.016)as measured by the Shiffman-Jarvik Withdrawal Scale (SJWS) subscale for craving.

